# Plastid gene expression is required for singlet oxygen-induced chloroplast degradation and cell death

**DOI:** 10.1101/2020.02.24.961144

**Authors:** Kamran Alamdari, Karen E. Fisher, Andrew B. Sinson, Joanne Chory, Jesse D. Woodson

## Abstract

Chloroplasts constantly experience photo-oxidative stress while performing photosynthesis. This is particularly true under abiotic stresses that lead to the accumulation of reactive oxygen species (ROS). While ROS leads to the oxidation of DNA, proteins, and lipids, it can also act as a signal to induce acclimation through chloroplast degradation, cell death, and nuclear gene expression. Although the mechanisms behind ROS signaling from chloroplasts remain mostly unknown, several genetic systems have been devised in the model plant *Arabidopsis* to understand their signaling properties. One system uses the *plastid ferrochelatase two* (*fc2*) mutant that conditionally accumulates the ROS singlet oxygen (^1^O_2_) leading to chloroplast degradation and eventually cell death. Here we have mapped three mutations that suppress chloroplast degradation in the *fc2* mutant and demonstrate that they affect two independent loci (*PPR30* and *mTERF9*) encoding chloroplast proteins predicted to be involved in post-transcriptional gene expression. Mutations in either gene were shown to lead to broadly reduced chloroplast gene expression, impaired chloroplast development, and reduced chloroplast stress signaling. In these mutants, however, ^1^O_2_ levels were uncoupled to chloroplast degradation suggesting that PPR30 and mTERF9 are involved in ROS signaling pathways. In the wild type background, *ppr30* and *mTERF9* mutants were also observed to be less susceptible to cell death induced by excess light stress. Together these results suggest that plastid gene expression (or the expression of specific plastid genes) is a necessary prerequisite for chloroplasts to activate ^1^O_2_ signaling pathways to induce chloroplast degradation and/or cell death.

**Significance summary:** Reactive oxygen species accumulate in the chloroplast (photosynthetic plastids) and signal for stress acclimation by inducing chloroplast degradation, cell death, and changes in nuclear gene expression. We have identified two chloroplast-localized proteins involved in gene regulation that are required to transmit these signals, suggesting that proper plastid gene expression and chloroplast development is necessary to activate chloroplast controlled cellular degradation and nuclear gene expression pathways.

## Introduction

Chloroplasts (differentiated plastids that perform photosynthesis) are required for the photoautotrophic lifestyle of plants, and by extension, most of life on the planet. While photosynthesis in these organelles is an essential process for most plants, it is also a problematic one. Photosynthesis itself involves highly reactive chemistry that leads to the formation of reactive oxygen species (ROS) such as singlet oxygen (^1^O_2_) at photosystem II (PSII) and superoxide and hydrogen peroxide (H_2_O_2_) at photosystem I (PSI) (Asada 2006). Although a certain amount of ROS can be produced even under ideal conditions, many common environmental stresses (e.g., excess light (Triantaphylides *et al.* 2008), drought (Chan *et al.* 2016), salinity (Suo *et al.* 2017), and pathogen attack (Lu and Yao 2018)) can drastically increase ROS production in the chloroplast.

Although chloroplasts still maintain small genomes containing ~100 genes encoding proteins, rRNA’s, and tRNA’s, the remaining ~3,000 proteins in chloroplasts are encoded in the nucleus (van Wijk and Baginsky 2011). Thus, chloroplasts need to be in constant communication with the rest of the cell to control their own proteome, ROS accumulation, and ROS damage under dynamic environments. This communication is achieved, in part, by the ROS themselves. Although ROS can lead to oxidative damage of proteins, lipids, and metabolites, they can also act as signaling molecules leading to chloroplast degradation, cell death, or changes in nuclear gene expression (Foyer 2018). Therefore, it appears chloroplast ROS can play two roles; as a stress signal to prepare a cell for oxidative damage and as information about the changing external environment to allow the cell (and plant) make the necessary biochemical and physiological changes to maintain efficient photosynthesis and to survive.

Exactly how ROS made in the chloroplast can have specific signaling abilities is still unclear. In *Arabidopsis*, it is known that ^1^O_2_ can signal independently of other chloroplast ROS such as H_2_O_2_ (op den Camp *et al.* 2003) and that ^1^O_2_ has the ability to initiate cellular degradation and nuclear gene reprogramming (Ramel *et al.* 2013, Wagner *et al.* 2004, Woodson 2019). These studies have mostly involved ^1^O_2_ accumulating mutants of *Arabidopsis* such as *plastid ferrochelatase II* (*fc2*), which conditionally accumulates the photosensitizer tetrapyrrole intermediate protoporphyrin IX (Proto) at dawn under diurnal light cycling conditions (Woodson *et al.* 2015). This accumulation of Proto in the light leads to a burst of chloroplast ^1^O_2_ that initiates chloroplast degradation and eventual cell death. A large forward genetic screen for suppressors of the *fc2* cell death phenotype identified 24 *ferrochelatase two suppressors* (*fts*) mutants affecting 17 loci. Some of these mutations could reverse the *fc2* chloroplast degradation and cell death phenotypes without affecting ^1^O_2_ accumulation. This suggested that such cellular degradation was not due to the toxicity of ^1^O_2_, but rather a genetic signal. One such mutant, *pub4-6* (*a.k.a. fts29*), was demonstrated to affect Plant U-Box 4, a cytoplasmic ubiquitin E3 ligase. This led to a model whereby cellular ubiquitination machinery is involved in targeting damaged chloroplasts for turnover (Woodson 2016). Such a chloroplast quality control mechanism may permit a cell to maintain a healthy population of chloroplasts under stressful conditions.

How ^1^O_2_ is able to initiate such a signal remains unknown. Due to its extremely short half-life (0.5–1 ms in cells (Ogilby 2010)), ^1^O_2_ is unlikely to travel further than a few hundred nanometers while chloroplasts are usually 5-10 microns across. Thus, the ^1^O_2_ produced within a particular chloroplast is unlikely to exit and must generate secondary signals, possibly in the form of oxidized proteins, lipids, or metabolites. Genetic studies have shown that multiple chloroplast ^1^O_2_ signaling pathways exist, possibly originating from distinct and specific sub-compartments of the chloroplast (Woodson 2019). Executor 1 (EX1) (Wagner *et al.* 2004) was the first chloroplast ^1^O_2_ signaling protein identified to be involved in transmitting ^1^O_2_ signals to the nucleus to modulate gene expression. However, it does not appear to be involved in chloroplast quality control as *ex1* mutations do not effect chloroplast degradation or nuclear gene expression in *fc2* mutants (Woodson *et al.* 2015). Therefore, to further understand the chloroplast quality control pathway and to identify additional genes involved, we have returned to the genetic *fc2-1* suppressor screen.

To bulk map the original *fts* mutations, the *fts* mutants were backcrossed to the *fc2-1* parent and the F2 plants exhibiting an *fts* phenotype were pooled for genomic DNA isolation, followed by library preparation, and next generation sequencing (Woodson *et al.* 2015). Shore-mapping analysis was used to identify the EMS-generated SNPs and their proximity to the causative allele (Schneeberger *et al.* 2009). This resulted in the rough-mapping of each mutation to a 1-3 mb region and the identity of the 10-30 SNPs in those regions. In this study, we continued with the analysis of three additional *fts* mutant lines, and demonstrate that they contain mutant alleles of *PPR30* (*fts3* and *fts38*) (a member of the pentatricopeptide repeat (PPR) containing protein family) or *mTERF9* (*fts32*) (a member of the “mitochondrial” transcription termination factor (mTERF) protein family).

Both PPR and mTERF proteins are predicted to be involved in post-transcriptional gene expression (Barkan and Small 2014, Kleine 2012). Although the chloroplast genome normally contains fewer than 100 genes encoding proteins, rRNA’s, and tRNA’s, their expression is essential for chloroplast development and photosynthesis (Yu *et al.* 2014). Chloroplasts encode one of their three RNA polymerases, the plastid encoded RNA polymerase (PEP), but gene expression is primarily controlled by nuclear-encoded factors (such as PPR and mTERF proteins) influencing post-transcriptional regulation (Barkan and Goldschmidt-Clermont 2000).

Here, we demonstrate that PPR30 and mTERF9 are chloroplast nucleoid-localized proteins and their absence leads to broadly impaired plastid gene expression and chloroplast development. Furthermore, mutations in *PPR30* and *mTERF9* block ^1^O_2_ signals that regulate chloroplast degradation, cell death, and retrograde signaling to the nucleus. Together our results suggest that chloroplast gene expression is a necessary prerequisite for transmitting the ^1^O_2_ signal to initiate chloroplast quality control pathways.

## Results

### Mapping of three *fts* alleles; *fts3*, *fts32*, and *fts38*

Previously, eight *fts* alleles were mapped and shown to represent four loci involved in chloroplast quality control (Woodson *et al.* 2015). To find additional components in this pathway, we chose three more *fts* mutants for further study (Table S1). As shown in Fig. 1a, while *fc2-1* mutants fail to green in 6h light/18h dark diurnal cycling conditions, *fc2-1 fts3*, *fc2-1 fts32*, and *fc2-1 fts38* survive. Under these conditions, cell death was suppressed in the double mutants as illustrated by reduced trypan blue staining (Fig. 1b). Like many of the other *fts* mutants isolated (Woodson *et al.* 2015), these three mutants also accumulated less chlorophyll than the *fc2-1* parent in constant light conditions (Fig. 1c) and had pale phenotypes (Fig. 1a).

**Figure 1.**
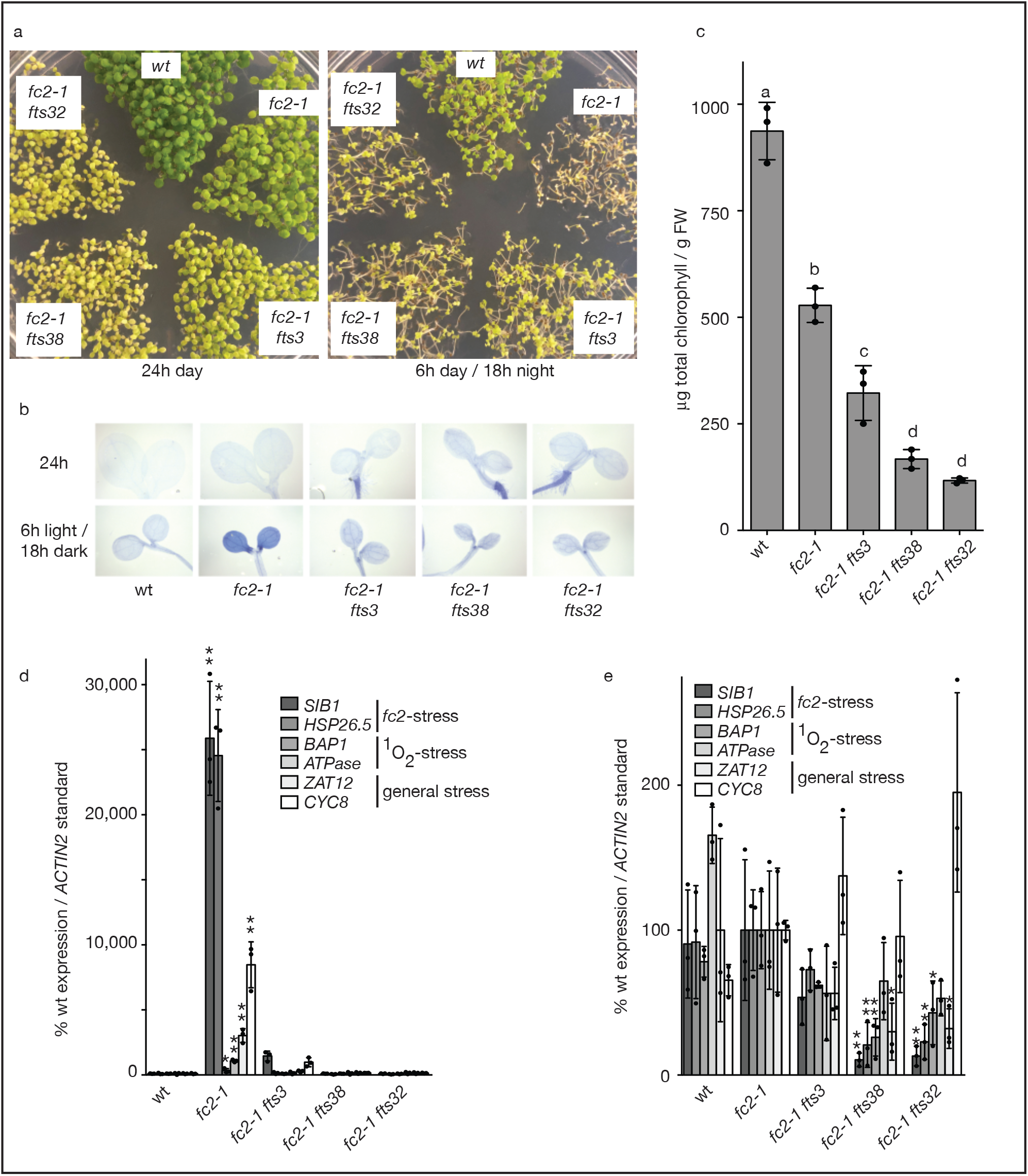
*fts3*, *38*, *32* are robust suppressors of the *fc2* cell death phenotype. *fc2-1 fts3, fc2-1 fts38, and fc2-1 fts32* mutants suppress the *fc2-1* cell death phenotype. **A)** *fc2-1 fts3, fc2-1 fts38, and fc2-1 fts32* were isolated based on their ability to survive under diurnal light cycling conditions. Shown are seven-day-old seedlings grown in constant (24h) light or diurnal cycling (6h days / 18h nights) light. **B)** Shown are representative seven-day-old seedlings stained with trypan blue. The deep dark color is indicative of cell death. **C)** Total chlorophyll levels of five-day-old seedlings grown in constant light. Shown are means of biological replicates (n = 3) +/− SD. The chlorophyll statistical analysis was performed by a one-way ANOVA and different letters above bars indicate significant differences determined by a Tukey-Kramer post-test (p value ≤ 0.05). RT-qPCR analysis of transcripts from 4 day old seedlings grown in **D)** 6 hours of light / 18 hours dark diurnal cycling and **E)** constant light conditions. Seedlings were harvested one hour post subjective dawn. Shown are means of biological replicates (n = 3) +/− SD. The RT-qPCR statistical analyses were performed by a one-way ANOVA followed by Dunnett’s multiple comparisons test with the wt (D) or *fc2-1* (E) sample. * and ** indicates an adjusted p value of ≤ 0.05 and ≤ 0.01, respectively. In all bar graphs, closed circles represent individual data points.

To determine if chloroplasts in these lines are transmitting retrograde signals to the nucleus in response to photo-oxidative stress, the expression of six marker genes for damaged chloroplasts (Woodson *et al.* 2015), chloroplast ^1^O_2_ (op den Camp *et al.* 2003), and general stress (Baruah *et al.* 2009) were monitored by RT-qPCR. We harvested four day old seedlings one hour after subjective dawn, which is when such stress genes are activated in *fc2-1* mutants (Woodson *et al.* 2015). In 6h light/18h dark diurnal cycling conditions, expression of all six genes were significantly increased in *fc2-1* seedlings compared to wt (Fig. 1d). In the double mutants, however, no single gene was significantly induced compared to wt. Next, we monitored the expression of the same genes in permissive constant light conditions when *fc2-1* mutants do not experience wholesale chloroplast degradation and cell death (Fig. 1e) (Woodson *et al.* 2015). Under these conditions, four of these marker genes were significantly repressed in *fc2-1 fts32* and *fts2-1 fts38* compared to *fc2-1.* Together, these results indicate that there was no initiation of chloroplast-to-nucleus (retrograde) stress signals in these double mutant lines.

Together, the phenotypes of these three suppressors were similar to the previously mapped *fts4*/*37*, *fts1*, and *fts8*/*16*/*24*/*30* mutations that are mutant alleles of *ppi1, ppi2*, and *chlH*, respectively (Woodson *et al.* 2015). PPI1 and PPI2 are plastid outer envelope-localized protein importers, while CHLH is an essential component of the Proto Mg-chelatase complex at the first committed step of the chlorophyll branch of the chloroplast-localized tetrapyrrole biosynthesis pathway. To investigate the possibility that *fts3*, *fts32*, and *fts38* also contained mutations affecting plastid protein import and/or tetrapyrrole synthesis, we used our sequencing data to determine if any of the ~1,000 SNP’s in each line is in the coding sequence of a gene involved in either process. Our cross-referencing of genes involved in tetrapyrrole biosynthesis (Tanaka *et al.* 2011) or plastid protein import (Jarvis and Lopez-Juez 2013) failed to find any putative mutant alleles in the three *fts* mutant lines. As such, we concluded that these three mutants may be affecting a different process related to chloroplast quality control.

The causative mutations in *fts3* and *fts38* had been mapped to the same region on the north arm of chromosome three (6.2-9.2 mb) suggesting that they may contain mutant alleles of the same gene (Woodson *et al.* 2015). A comparison of genes with SNPs in their coding regions identified *AT3g23020* as the only gene mutated in both *fts3* and *fts38* (Fig. 2a). *AT3g23020* is predicted to encode PPR30, belonging to the PPR family of organelle proteins that bind specific RNA transcripts and can regulate their expression by influencing stability, translation, splicing, and editing (Barkan and Small 2014). *fts3* and *ft38* have mutations leading to the amino acid substitutions D765N and G821R, respectively. A protein alignment suggests that these residues are conserved among dicots, while D765 and G821R reside within PPR motif 16 or just outside PPR motif 17, respectively (Fig. 2b). Interestingly, null mutations in *PPR30* have been observed to be embryo lethal, suggesting that *PPR30* encodes an essential gene (Savage *et al.* 2013). Therefore, it is possible that *fts3* and *fts38* encode weak loss-of-function *ppr30* alleles.

**Figure 2.**
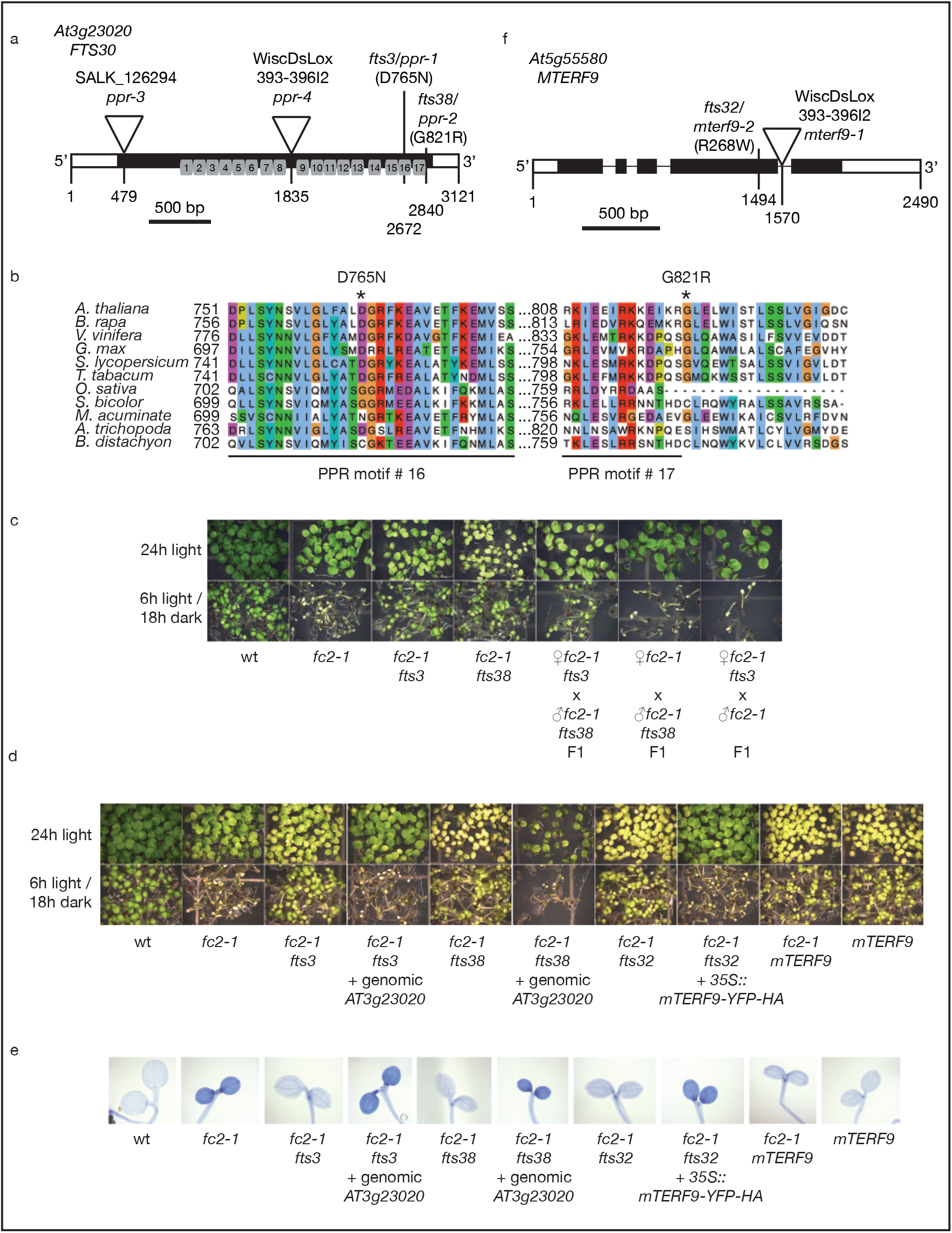
Mapping of *fts3*, *fts32*, and *fts38*. *fts3*, *38*, and *32* were mapped to two independent loci encoding proteins putatively involved in chloroplast gene expression. **A)** A schematic of the *PPR30* (*At3g23020*) gene in *Arabidopsis*. Shown is the position of the EMS generated *fts3* and *fts38* alleles identified in the *fc2-1* suppressor screen and two previously described T-DNA insertion lines (Savage *et al.* 2013). Predicted PPR motifs 1-18 are indicated under the FTS3 schematic. **B)** Alignment of FTS3 orthologs in various land plants. Numbers on the left hand side indicate amino acid positions. The two amino acids substituted in the *fts3* and *fts38* mutants are indicated with asterisks. **C** and **D)** Representative images of six-day-old seedlings grown in constant (24h) light or 6h light / 18h dark diurnal cycling conditions. **He)** Shown are representative six-day-old seedlings from **D** stained with trypan blue. The deep dark color is indicative of cell death. **F)** A schematic of the *FTS32/mTERF9* (*At5g55580*) gene in *Arabidopsis.* Shown is the position of the EMS generated *fts32* allele identified in the *fc2-1* suppressor screen. Also shown are the positions of the previously described T-DNA insertion line (Robles *et al.* 2015).

To confirm if *fts3* and *fts38* are allelic, we performed crosses between the two lines and tested the *fts* phenotype of the resulting F1 generation. In agreement with *fts3* and *fts38* being allelic, F1 seedlings from the cross were similar to both *fc2-1 fts3* and *fc2-1 fts38* and survived 6h light/18h dark diurnal cycling conditions (Fig. 2c). On the other hand, *fc2-1 fts3* or *fc2-1 fts38* backcrossed to *fc2-1* resulted in F1 seedlings with an *fc2-1* phenotype (seedlings failed to green in 6h light/18h dark diurnal cycling conditions) confirming that both mutations are recessive. A genomic fragment containing the *PPR30* coding, promoter, and terminator sequences was cloned into a binary vector and transformed into *fc2-1 fts3* and *fc2-1 fts38.* In the T3 generation of both lines, this wt copy of *PPR30 r*estored the parent *fc2-1* phenotype (the seedlings failed to green in 6h light/18h dark diurnal cycling conditions (Fig. 2d) and cell death was observed through trypan blue staining (Figs. 2e, S1a and b)). Together, these results clearly indicate that the mutations in *AT3g23020* were responsible for the *fts* phenotypes of *fc2-1 fts3* and *fc2-1 fts38.* Furthermore, the more pronounced chlorophyll deficiency (Fig. 1c) and repression of nuclear stress genes (Fig. 1d and e) suggests that *fts38* is the stronger allele of the two.

As *fts3* and *fts38* may affect gene expression in the chloroplast, we scanned the mapped region of *fc2-1 fts32* (chromosome five 22.2mb – 24.9mb) for mutations in genes encoding chloroplast proteins. In this region, we found that *At5g55580*/*mTERF9* contained a mutation leading to a premature stop codon at residue R368 (Fig. 2f). *mTERF9* has previously been shown to encode a protein that is involved in chloroplast gene expression (Robles *et al.* 2015). To determine if this mutation was responsible for the *fts32* phenotype, we took two approaches. First, we transformed *fc2-1 fts32* mutant plants with a *35S::mTERF9-YFP* construct that produces a wt mTERF9 protein with a C-terminal YFP tag. This construct was able to fully complement the *fts32* phenotype leading to cell death (Figs. 2e, S1a and b) and the failure to green under 6h light/18h dark diurnal cycling conditions (Fig. 2d). At the same time, we obtained a second allele of *mTERF9* containing a T-DNA in its fourth intron leading to a loss-of-function phenotype (Fig. 2f) (Robles *et al.* 2015). This allele was crossed into the *fc2-1* background and tested for an *fts* phenotype. As expected, this allele phenocopied the *fts32* mutation and *fc2-1 mterf9* was able to green under 6h light/18h dark diurnal cycling conditions and avoid cell death (Figs. 2d and e, S1a and b). Together, these results confirm that the *mTERF9* nonsense mutation in *fc2-1 fts32* is the causal mutation leading to suppression of *fc2* phenotypes.

### AT3g23020 and mTERF9 are localized to chloroplasts

To determine the subcellular localization of PPR30 and mTERF9, we transiently produced these proteins with C-terminal YFP tags in *Nicotine benthamiana* leaves and monitored YFP localization using a laser scanning confocal microscope. As shown in Fig. S2, PPR30-YFP and mTERF9-YFP clearly overlapped with chlorophyll autofluorecence in the leaf mesophyll cells indicating that these two proteins are localized to chloroplasts. This result confirmed an earlier report that mTERF9 is exclusively localized to chloroplasts and not mitochondria (Babiychuk *et al.* 2011). Interestingly, with both constructs, a speckle-like pattern of YFP signal was also observed in these chloroplasts. Such a pattern indicates a localization to chromatin-dense nucleoid complexes within these organelles and is consistent with a role in plastid gene expression (Yagi and Shiina 2014).

### *fc2-1 fts32* and *fc2-1 fts38* mutants have delayed chloroplast development

Because the *fts3*/*38* and *fts32* mutations lead to reduced chlorophyll accumulation in the *fc2-1* background (Fig. 1c), we hypothesized that these mutants may have impaired chloroplast function or development. One hallmark of such defects is the down-regulation of photosynthesis associated nuclear genes (PhANGs) by retrograde signals from developing chloroplasts (Chan *et al.* 2015). Using RT-qPCR, we monitored the expression of four PhANGs (*LHCB1.2*, *LHCB1.2*, *CA1*, and *RBCS2b*) in 4-day-old seedlings grown in constant light. Consistent with these lines having delayed chloroplast development, the expression of at least two PhANGs were significantly down-regulated compared to *fc2-1* in each of the three double mutants. In the case of *fc2-1 fts38*, all four genes were significantly repressed (Fig 3a), further confirming that *fts38* is the stronger *ppr30* allele.

**Figure 3.**
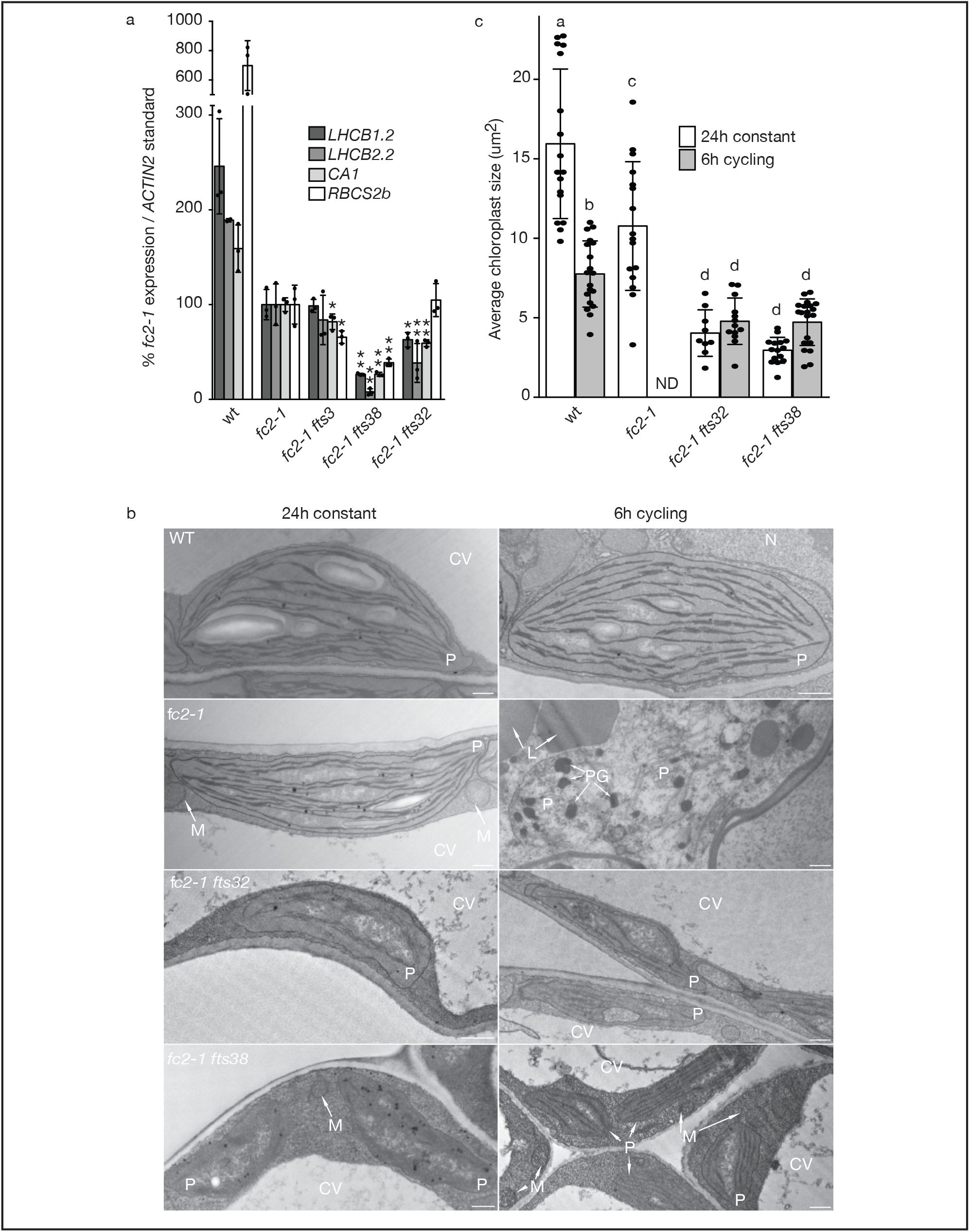
*fc2-1 fts32* and *fc2-1 fts38* have delayed chloroplast development. **A)** RT-qPCR analysis of transcripts from 4 day old seedlings grown in constant light. Shown are means of biological replicates (n=3) +/− SD. Statistical analysis was performed by a one-way ANOVA followed by Dunnett’s multiple comparisons test with the *fc2-1* sample. * and ** indicate an adjusted p value of ≤ 0.05 and ≤ 0.01, respectively. **B)** Shown are representative TEM micrographs of chloroplasts in cotyledon mesophyll cells of four day-old seedlings grown in 24h constant light or 6h light/18h dark diurnal cycling conditions one hour after subjective dawn. Plastids (P), Nuclei (N), central vacuoles, (CV), lipid bodies (L), plastoglobules (PG), and mitochondria (M) are noted. Scale bars = 500 nm. **C)** Shown are means of the average size (um^2)^ of chloroplasts in b (+/− SD) (n ≥ 9 cells). Statistical analysis was performed by a one-way ANOVA and different letters above bars indicate significant differences determined by a Tukey-Kramer post-test (p value ≤ 0.05). In all bar graphs, closed circles represent individual data points.

Next, we attempted to visualize the ultrastructure of cotyledon mesophyll chloroplasts. Four day old seedlings grown in both constant light and 6h light/18h dark diurnal cycling conditions were fixed and imaged by transmission electron microscopy (TEM). As was expected from the phenotypes, *fc2-1* chloroplasts were degraded under 6h/day light diurnal cycling conditions (Fig. 3b). Such chloroplasts have poorly defined envelopes, a loss of organized thylakoid structures, and large plastoglobules (Woodson *et al.* 2015). Conversely, chloroplasts in *fc2-1 fts32*, *fc2-1 fts38*, and wt were intact. Interestingly, under constant light conditions, *fc2-1 fts32* and *fc2-1 fts38* chloroplasts were also irregularly shaped, contained fewer internal membranes, were significantly smaller, and constituted a smaller portion of the cell compared to wt and *fc2-1* (Figs. 3c and S3). Unlike wt chloroplasts, a short-day length did not appear to reduce chloroplast size or development in *fc2-1 fts32* and *fc2-1 fts38* mutants. Together, these results indicate that mutations in *mTERF9* and *PPR30* lead to chloroplasts that avoid degradation, are insensitive to light shifts, but are impaired in their development.

### *fts3*, *fts32*, and *fts38* are defective in plastid gene expression

As PPR30 and mTERF9 are chloroplast-localized proteins with predicted roles in gene expression, we tested if the *fc2-1 fts3*, *fc2-1 fts32*, or *fc2-1 fts38* mutants have altered levels of plastid-encoded transcripts. RT-qPCR analysis was used to assess the steady state levels of seven transcripts primarily transcribed by the plastid-encoded polymerase (PEP) (*spa, psbB, psaJ, rbcL, trnEYD*), the nuclear-encoded polymerase (NEP) (*rpoB*), or both (*clpP*) (Yagi and Shiina 2014). Unlike the other protein-encoding genes, *trnEYD* is a polycistronic transcript encoding the tRNA species for glutamate, tyrosine, and aspartate and is primarily regulated by the nuclear-encoded PEP subunit Sigma Factor 2 (SIG2) (Hanaoka *et al.* 2003). Glutamyl-tRNA is of special interest, as it is the starting precursor to tetrapyrrole (and chlorophyll) biosynthesis in addition to protein synthesis (Tanaka *et al.* 2011). Compared to *fc2-1*, the levels of two or four PEP transcripts were significantly reduced in *fc2-1 fts3* and *fc2-1 fts32*, respectively (Fig. 4). In the case of *fc2-1 fts38*, all five transcripts were significantly reduced including *trnEYD*, further confirming that *fts38* is the stronger *ppr30* allele. Unlike the PEP transcripts, the NEP transcripts were unresponsive or increased in the double mutants compared to *fc2-1*. Together, these results suggest that the *fts3/38* and *fts32* mutations lead to a broad defect in PEP-directed gene expression.

**Figure 4.**
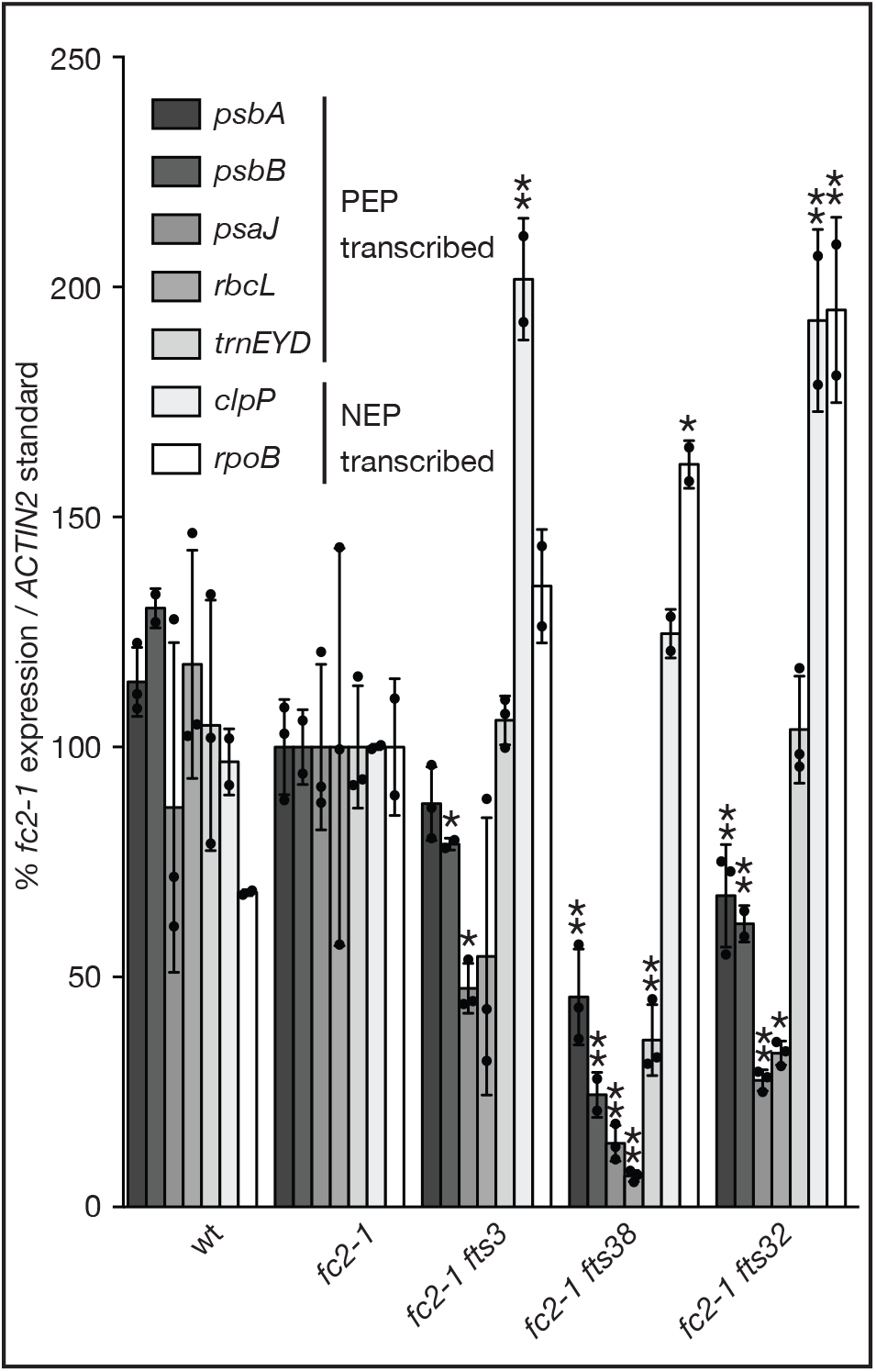
*fc2-1 fts3*, *fc2-1 fts38*, *fc2-1 fts32* mutants have impaired plastid gene expression. Shown are means (+/− SD) of expression levels of plastid transcripts from 4 day old seedlings grown in constant light as measured by RT-qPCR (n ≥ 2 biological replicates). Closed circles represent individual data points. Statistical analysis was performed by a one-way ANOVA followed by Dunnett’s multiple comparisons test with the *fc2-1* sample. * and ** indicate an adjusted p value of ≤ 0.05 and ≤ 0.01, respectively.

### *fts3*, *fts32*, and *fts38* have different effects on tetrapyrrole synthesis and singlet oxygen levels

In *fc2-1* mutants, ^1^O_2_ production is the result of the accumulation of Proto or other free tetrapyrroles (Scharfenberg *et al.* 2015, Woodson *et al.* 2015). Because tetrapyrrole synthesis is dependent on mature transcripts of *trnE*, we hypothesized that the *fts3/38* or *fts32* mutants could be indirectly reducing tetrapyrrole synthesis through reduced expression of *trnE* or by affecting the RNA’s maturation and processing. To this end, we tested the ability of the mutant lines to synthesize 5-aminolevulinic acid (ALA), the first committed step of the pathway. In six day-old seedlings grown in constant light, only the *fc2-1 fts38* mutant had reduced ALA synthesis compared to *fc2-1*. This suggested that early steps in tetrapyrrole synthesis had been impaired by the *fts38* mutation (Fig. 5a) possibly due to reduced *trnEYD* transcript. To determine if later steps of tetrapyrrole synthesis may be affected in these mutants, we next measured the accumulation of protochlorophyllide (Pchlide, a precursor of chlorophyll A) in etiolated (dark grown) seedlings. As previously shown (Scharfenberg *et al.* 2015, Woodson *et al.* 2015), the *fc2-1* mutant accumulated ~2.5-fold more Pchlide than wt (Fig. 5b). However, all three *fts* mutations reduced Pchlide back down to wt levels, suggesting that the Mg-branch of the tetrapyrrole pathway may be inhibited by the *fts3* and *fts32* mutations.

**Figure 5.**
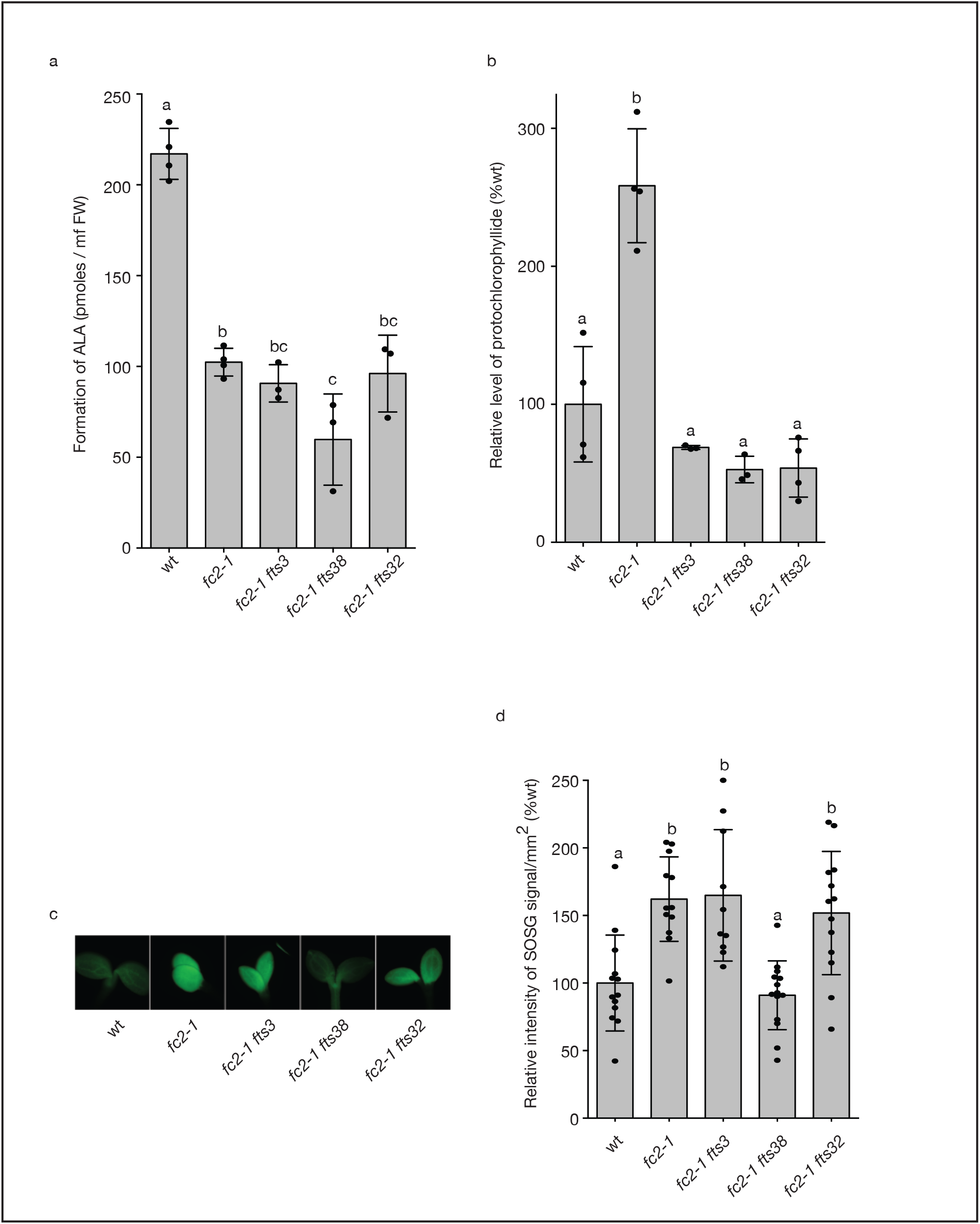
*fc2-1 fts3*, *fc2-1 fts38*, *fc2-1 fts32* mutants have impaired tetrapyrrole synthesis. **A)** ALA synthesis rates of six-day-old seedlings grown in constant light. Shown are means of biological replicate measurements (n ≥ 3) +/− SD. **B)** Protochlorophyllide levels of five-day-old seedlings grown in constant dark. Shown are means of biological replicate measurements (n ≥ 3) +/− SD. **C** and **D)** SOSG fluorescence in four day-old seedlings grown in diurnal cycling light (6 hours light/18h dark). Shown are representative seedlings and means of average intensities of SOSG signal across one cotyledon per seedling (n ≥ 10) +/− SD. Statistical analysis was performed by a one-way ANOVA and different letters above bars indicate significant differences determined by a Tukey-Kramer post-test (p value ≤ 0.05). In all bar graphs, closed circles represent individual data points

Next, we tested if reduced tetrapyrrole synthesis or plastid gene expression was affecting ^1^O_2_ production. The fluorescent ^1^O_2_ marker dye SOSG was infiltrated into four day-old seedlings grown under 6h light/18h dark diurnal cycling conditions and seedlings were monitored for fluorescence and the presence of ^1^O_2_ two hours after dawn (when ^1^O_2_ can be visualized in *fc2-1* seedlings (Woodson *et al.* 2015)). As expected, *fc2-1* mutants accumulated significantly higher levels of ^1^O_2_ than wt (Figs. 5c and d). Surprisingly, both *fc2-1 fts3* and *fc2-1 fts32* retained high levels of ^1^O_2_ similar to *fc2-1*, suggesting that *fts3* and *fts32* have uncoupled ^1^O_2_ production and chloroplast degradation in the *fc2-1* mutant. Conversely, *fc2-1 fts38* mutants had reduced ^1^O_2_ production similar to wt levels. This may be due to the relatively stronger phenotype of *fts38* leading to reduced *trnE* transcript levels and ALA synthesis.

### *fts3*, *fts32*, and *fts38* increase resistance to excess light

The previous results suggested that *fc2-1 fts3* and *fc2-1 fts32* still produce a significant amount of ^1^O_2_, but their chloroplasts do not suffer photo-oxidative damage and/or signal for chloroplast degradation. To further test the possibility that these mutants are resistant to ^1^O_2_, we outcrossed the double mutants to wt plants to isolate *fts3*, *fts32*, and *fts38* in wt backgrounds allowing us to test chloroplast stress in the absence of the *fc2-1* mutation. Plants were first grown for three weeks under mild light conditions of 100 μmol m^−2^ sec^−1^ white light (16h light/day) at 21°C. Individual leaves were then floated on liquid media and shifted to 650 μmol m^−2^ sec^−1^ white light at 4°C. The combination of excess light and low temperatures has previously been shown to lead to ^1^O_2_–induced cell death (Kim *et al.* 2012a, Meskauskiene *et al.* 2009, Triantaphylides *et al.* 2008). Over the course of 50h, leaves were stained for cell death with trypan blue. While wt leaves began to experience cell death after 18 hours, *fts3, fts38, fts32*, and *mTERF9* T-DNA mutant plants all suffered significantly less ^1^O_2_–induced cell death up through 50 hours of excess light (Figs. 6a and b). Exposing leaves to 4°C in dim light for a similar period of time did not induce cell death (except in the *mTERF9* T-DNA mutant, which exhibited a mild (but statistically significant) increase in trypan blue staining), suggesting that cell death in this experiment was largely dependent on photo-oxidative damage caused by excess light (Figs. S4a and b).

**Figure 6.**
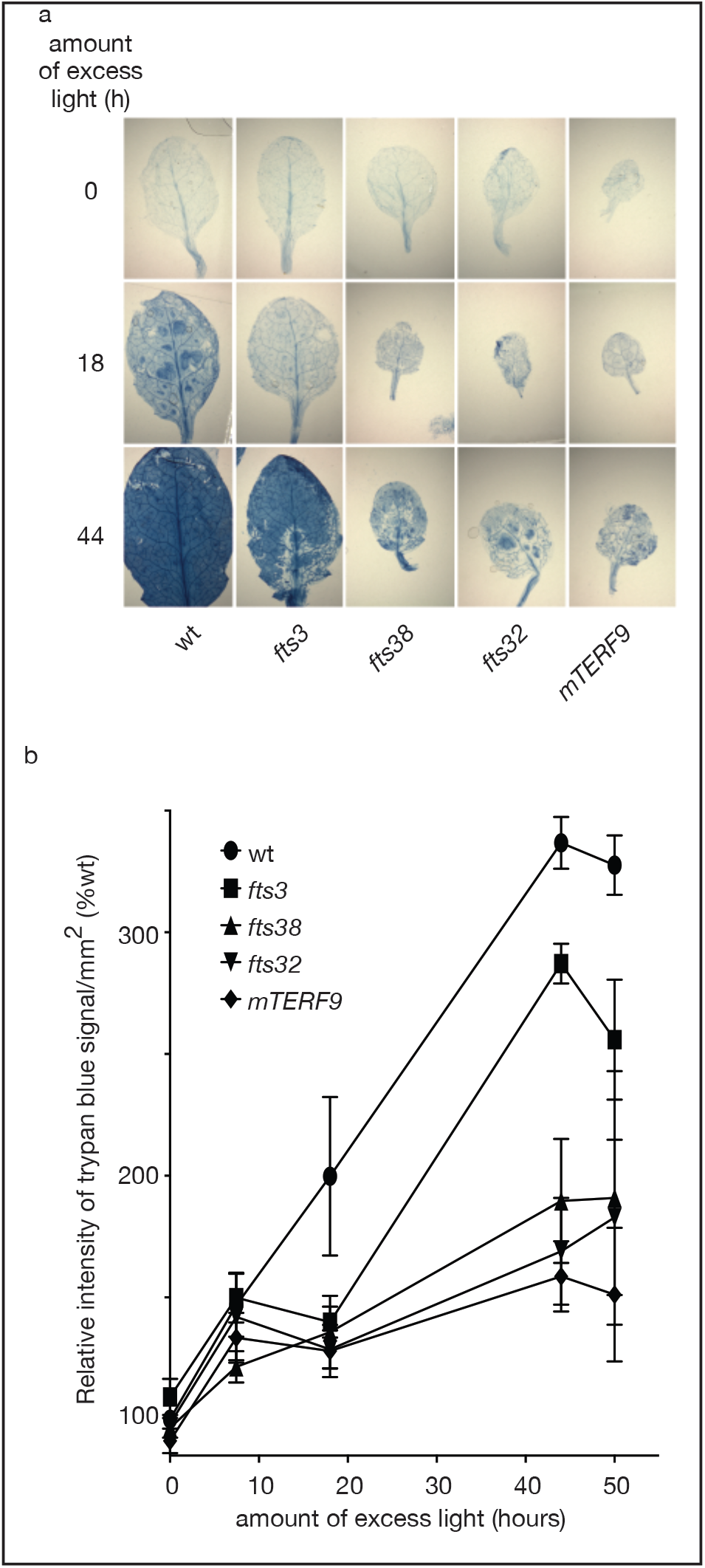
*ppr30* and *mTERF9* mutants are resistant to excess light stress. Single leaves of *ppr30* and *mTERF9* mutants were exposed to excess light (650 μmol m^−2^ sec^−1)^ and cold temperatures (4°C) for various lengths of time. **A)** Shown are representative images of these leaves stained by trypan blue. The deep blue color is indicative of cell death. **B)** Shown are the means of average intensities of the trypan blue signal/mm^2^ across the entire leaf (n = 6) +/− SD.

### *sig2*, *sig6, and gun1* do not lead to *fts* phenotypes

The conclusion that mutations in *ppr30* and *mTERF9* may lead to ^1^O_2_ resistance by reducing plastid gene expression suggests that any mutation negatively affecting plastid gene expression may also suppress *fc2-1* phenotypes in diurnal cycling light. One well-characterized mutation that leads to broadly impaired plastid gene expression is *sig2* (Hanaoka *et al.* 2003, Woodson *et al.* 2013), which has reduced expression of *psaJ, psaA, psaB, psbD, psbN* (Woodson *et al.* 2013) and tRNAs (trnE-UUC and trnV-UAC) (Hanaoka *et al.* 2003). This reduced *trnE* expression is likely responsible for the reduced ALA and tetrapyrrole (heme, Pchlide, and chlorophyll) synthesis in the *sig2* mutant (Woodson *et al.* 2013). It does not appear that *sig2* mutants were isolated in our original screen (Woodson *et al.* 2015). Of the remaining unmapped mutations, only *fts2*, *fts13*, and *fts40* were mapped near the location of *SIG2* (*At1g08540*). However, no mutation was found within ~26 kb of *SIG2* in these three mutants. Therefore, to test the possibility that *sig2* mutations may also lead to resistance to ^1^O_2_, we attempted to generate *fc2-1 sig2-2* double mutants by crossing. However, plants homozygous for both mutations were unable to be recovered on soil as they bleached and died before flowering. Therefore, we used a line that was homozygous for *sig2-2* and segregating for the *fc2-1* T-DNA allele. As shown in Fig. 7a, the *fc2-1* cell death phenotype was observed to be segregating in this background in a 1:3 ratio (X^2^ value = 0.214, p value 0.643, n = 224 seedlings) when seedlings were grown under 6h light / 18h dark diurnal cycling conditions. This ratio was consistent with these dead seedlings being *fc2 sig2-2* double mutants. Trypan blue staining confirmed these phenotypes and showed that the *fc2-1 sig2-2* mutants experienced more cell death than wt or *fc2-1 fts3* (Fig. 7b and c). Therefore, contrary to the other plastid gene expression mutants, the *sig2* mutation did not allow greening in diurnal light cycling conditions or suppress cell death (Figs. 7a, b, and c).

**Figure 7.**
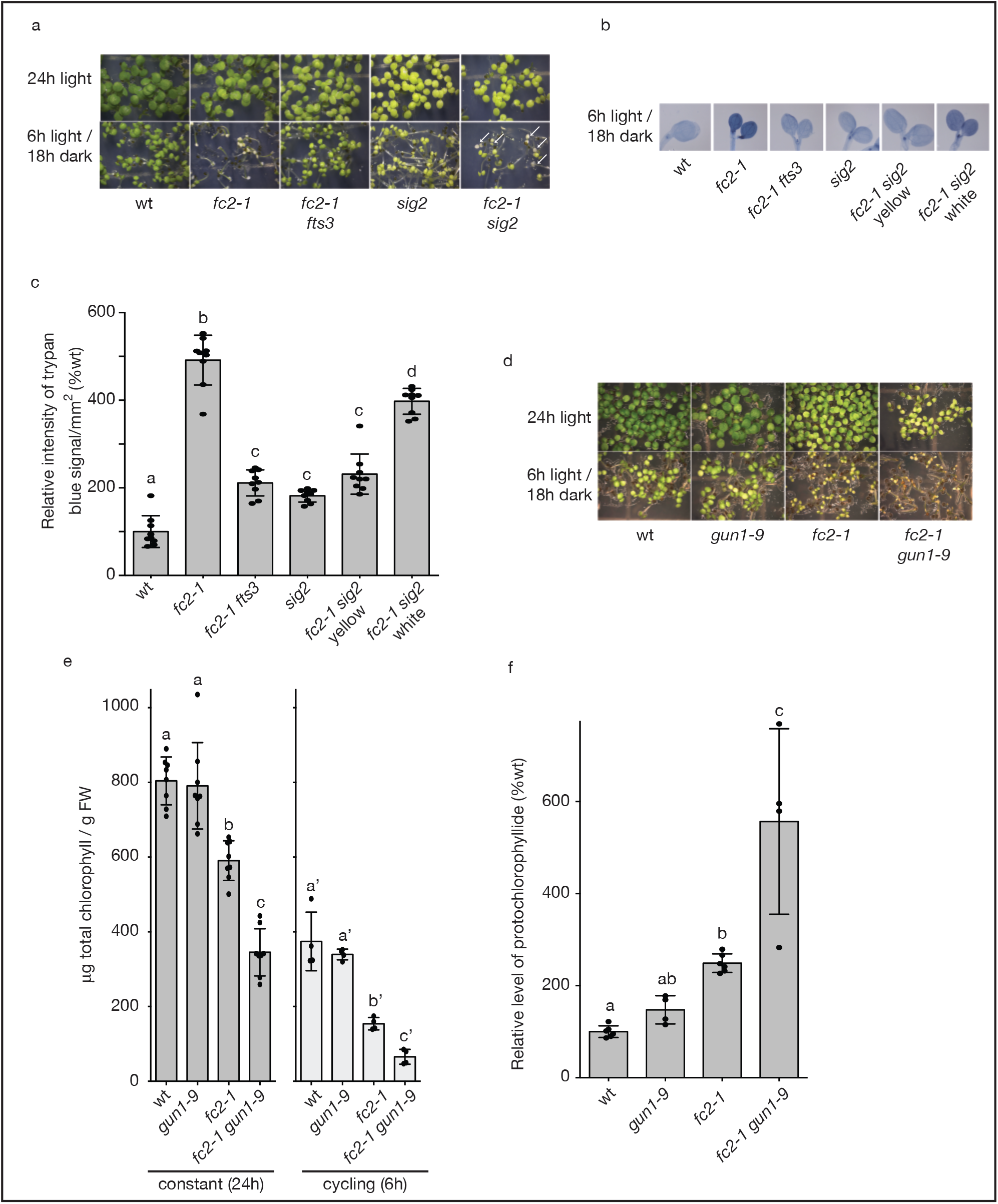
*fts3*, *38*, *32* mutants have specific defects in gene expression. *sig2* and *gun1* mutations do not suppress the *fc2-1* cell death phenotype under diurnal light cycling conditions. **A)** Shown are six-day-old seedlings grown in constant (24h) light or diurnal cycling (6h days / 18h nights) light. **B)** Shown are representative six-day-old seedlings grown in diurnal cycling conditions and stained with trypan blue. The deep dark color is indicative of cell death. **C)** Quantification of the trypan blue staining (relative intensity/mm^2)^ (means of nine seedlings +/− SD). **D)** Shown are four-day-old seedlings grown in constant light or diurnal cycling light. **E)** Total chlorophyll levels of same seedlings. Shown are means of biological replicates (n = 8, 24h samples; 4, 6h samples) +/− SD. **F)** Pchlide levels of five-day-old seedlings grown in constant dark. Shown are means of biological replicates (n ≥ 4) +/− SD. Statistical analysis was performed by a one-way ANOVA and different letters above bars indicate significant differences determined by a Tukey-Kramer post-test (p value ≤ 0.05). In all bar graphs, closed circles represent individual data points.

Like PPR30, GUN1 is another chloroplast-localized PPR protein that has been implicated in chloroplast stress and signaling (Koussevitzky *et al.* 2007). When chloroplast development has been blocked, *gun1* mutants can also have reduced chloroplast gene expression (Woodson *et al.* 2013). As such, we tested the possibility that *gun1* mutants may also suppress the *fc2-1* phenotype. A *fc2-1 gun1-9* double mutant was generated by crossing and then grown in 24h or 6h light/18h dark diurnal cycling conditions. Instead of suppressing the *fc2-1* phenotype, *gun1-9* appeared to enhance it; *fc2-1 gun1-9* seedlings were much paler than *fc2-1* mutants in 24h light conditions and appeared to suffer more photo-oxidative damage in 6h light / 18h dark diurnal cycling conditions as shown by reduced chlorophyll accumulation (Figs. 7d and e). As recent reports have suggested that GUN1 may be involved in tetrapyrrole synthesis, we measured the levels of Pchlide in five day-old etiolated seedlings (Fig. 7f). Surprisingly, *fc2-1 gun1-9* mutants had significantly elevated Pchlide compared to *fc2-1*. Together these results suggest that by increasing tetrapyrrole synthesis, *gun1-9* mutants may also increase the sensitivity of *fc2-1* to diurnal cycling light.

## Discussion

During photosynthesis, chloroplasts constantly experience stress in the form of ROS, which has the ability to cause extensive oxidative damage to the organelle and the cell. Although it is known that such ROS can lead to signals that control chloroplast degradation, cell death, and nuclear gene expression, how ROS initiates these signals and how these signals are propagated remains unclear. This is due, in part, to the complexity of different ROS being produced under natural stress conditions (e.g., excess light, drought) that likely lead to multiple signaling networks being activated in a single cell (Chan *et al.* 2015, Woodson 2019). However, the use of *Arabidopsis* mutants that specifically produce only one ROS in the chloroplast has allowed for the study of these pathways in isolation. The study of one such mutant, *fc2-1*, has revealed a ^1^O_2_ pathway that leads to the degradation of severely damaged chloroplasts. The purpose of such a chloroplast quality control pathway may be to ensure a healthy population of photosynthesizing chloroplasts in a given cell. Although it is hypothesized that ^1^O_2_ accumulation leads to the ubiquitination of chloroplast envelope proteins to “mark” a chloroplast for degradation, how this signal is initiated within the chloroplast is still not known.

To address this and to identify such signaling factors, here we have mapped three additional *fts* mutations that suppress the *fc2-1* cell death phenotype under diurnal light cycling conditions. Two of these alleles (*fts3* (*fts30-1*) and *fts38* (*fts30-2*)) have point mutations in *At3g20320* (*FTS30*), which encodes a chloroplast-localized PPR protein. The PPR proteins are a greatly expanded group of proteins in land plants, with most species having over 400 members (Small and Peeters 2000). In most cases these proteins are targeted to organelles (both plastids and mitochondria) where they play a role in almost every step of gene expression including transcription, RNA stabilization, RNA cleavage, RNA splicing, RNA editing, and translation (Barkan and Small 2014). PPR30 specifically belongs to the P type PPR protein that contains the archetypal 35 amino acid P motif (Small and Peeters 2000). This class of protein is usually involved in the stabilization of specific RNA’s, but it can also promote cleavage, translation, and in rare cases, editing (Doniwa *et al.* 2010). The exact role of PPR30 is still unknown, but here we show that *ppr30* mutations reduce the accumulation of several PEP-dependent transcripts and lead to impaired chloroplast development. However, a previous report has shown that null alleles of *PPR30* are embryo lethal (Savage *et al.* 2013) suggesting a role beyond photosynthesis. Therefore, PPR30 may be involved in the more general process of plastid translation (possibly through the expression of an rRNA transcript), thus affecting PEP formation indirectly. Consistent with this hypothesis, the *Zea mays* ortholog of PPR30 shows the strongest expression near the base of the maize leaf, suggesting it has a role in the build-up of plastid gene expression machinery in the very early stages of chloroplast development (Li *et al.* 2010).

Another PPR protein known to have an important role in chloroplast signaling is GUN1, which is necessary for the coordination of photosynthetic gene expression between the plastid and nuclear genomes (Koussevitzky *et al.* 2007). Interestingly, GUN1 may be an unusual case as it has not yet been shown to target a specific RNA transcript (Pesaresi and Kim 2019). Instead, GUN1 has been proposed to bind to several different proteins in the chloroplast including import-related chaperone cpHSC70-1 (Wu *et al.* 2019), the MORF2 editing protein (Zhao *et al.* 2019), and Plastid Ferrochelatase 1 (FC1) (Shimizu *et al.* 2019). Here we establish that a *gun1* mutation clearly does not suppress *fc2-1* or lead to an *fts* phenotype. Instead, *fc2-1 gun1-9* mutants appear to suffer enhanced photo-damage. This appears to be due to an increase of tetrapyrrole synthesis in this mutant and is consistent with a recent report demonstrating that GUN1 plays a role in tetrapyrrole metabolism by directly binding to tetrapyrroles such as Proto and by stimulating FC1 activity (Shimizu *et al.* 2019). As such, by modulating tetrapyrrole synthesis or by sequestering Proto, GUN1 may play an indirect role in ^1^O_2_ signaling from the chloroplast.

A third mutant, *fts32*, was shown to affect the function of mTERF9 by introducing a premature stop codon in its coding sequence. Like PPR proteins, mTERFs are also part of a large family of proteins with roles in post-transcriptional gene expression in organelles. Although most are uncharacterized, mTERF proteins appear to regulate transcriptional termination and RNA processing (Kleine 2012). Here we demonstrate that mTERF9 localizes to chloroplast nucleoids, which is in agreement with a previous nucleoid proteomic study (Majeran *et al.* 2012). Also like PPR30, loss of mTERF9 using the *fts32* allele leads to a broad reduction of PEP-dependent gene expression and impaired chloroplast development. Robles *et al.* (2015) also reported that the null mTERF9 T-DNA line leads to reduced *psbA* and *psbD* transcript levels and changes in chloroplast ultrastructure. Unlike PPR30, however, mTERF9 is dispensable for growth as the same mutant is viable without any supplemental carbon source. This may suggest that mTERF9 directs the expression of a different component in the chloroplast or is a partially redundant protein. However, the target gene of mTERF9 is still unknown.

The commonality behind the *ppr30* and *mterf9* mutants is that they all broadly reduce PEP-dependent gene expression in the chloroplast. As such, how can the reduction of plastid gene expression lead to an *fts* phenotype and suppress chloroplast degradation in the *fc2-1* mutant? We tested the possibility that these mutants were reducing available transcripts of *trnE*, which would limit the synthesis of tetrapyrroles and thus, ^1^O_2_. However, the reduction of *trnE* and ALA was not the common factor among the mutants. Only the stronger allele of *ppr30* (*fts38*) led to a mild reduction of *trnE* transcripts, ALA synthesis, and subsequent ^1^O_2_ levels. Although later steps in tetrapyrrole synthesis were affected by the *mterf9* and weak *ppr30* (*fts3*) mutations (as evidenced by reduced steady-state Pchlide levels in etiolated seedlings), this was not sufficient to reduce ^1^O_2_ levels under diurnal light cycling. This suggests that by eliminating mTERF9 or mildly reducing PPR30 content, chloroplast degradation and cell death can be uncoupled with ^1^O_2_ accumulation in *fc2* mutants.

This is reminiscent of the mutants *soldat8* (Coll *et al.* 2009) and *soldat10* (Meskauskiene *et al.* 2009), that have mutations in *SIGMA6* and *mTERF1*, respectively. Both were found as suppressors of ^1^O_2_-induced cell death in the *fluorescent in blue* light (*flu*) mutant that over-accumulates the photosensitizer tetrapyrrole Pchlide in the dark. Their mode of suppression, however, appears to involve disrupting chloroplast homeostasis and promoting a constitutive stress response in the nucleus through retrograde signals. Our RT-qPCR analysis of stress-related transcripts in *fc2-1 ppr30* and *fc2-1 mterf9* mutants showed that such transcripts are clearly not upregulated under permissive constant light conditions. Furthermore, unlike the *soldat* mutants, ^1^O_2_-marker genes were also not upregulated under stress conditions (diurnal cycling light in this report). Together this suggests that the *ppr30* and *mterf9* mutations are blocking the ability of ^1^O_2_ to signal outside the chloroplast. While the short half-life (~1 usec (Ogilby 2010)) of ^1^O_2_ makes it an unlikely signaling molecule, it is possible that an oxidation product (a chloroplast lipid, protein, or metabolite) may be the true signal that leads to chloroplast degradation and retrograde signaling, possibly by affecting chloroplast ubiquitination or a cytoplasmic signaling cascade. Such a hypothetical chloroplast metabolite or protein may then be less abundant in *ppr30* and *mterf9* mutants due to decreased gene expression and delayed chloroplast development. One protein known to transmit a ^1^O_2_ signal this way is EX1, whose oxidation by ^1^O_2_ and subsequent degradation by FtsH2 (Dogra *et al.* 2017, Dogra *et al.* 2019) initiates a retrograde signal to control nuclear gene expression. However, *ex1* mutations do not suppress *fc2* mutants, suggesting that these pathways do not directly interact (Woodson *et al.* 2015) and that PPR30 and mTERF9 likely affect a different signaling factor.

Rose Bengal (a photo sensitizer dye that leads to ^1^O_2_ production) has previously been shown initiate cell death in *Arabidopsis* cell cultures, but only those with mature chloroplasts (Gutierrez *et al.* 2014). Along with the work reported here, these results demonstrate a strong link between the developmental stage of a chloroplast (or plastid) and its ability to respond to ^1^O_2_. However, it does not appear that generally reducing plastid gene expression and development is always sufficient to lead to an *fts* phenotype and block ^1^O_2_ signaling. The *sig2* mutation has previously been shown to lead to a broad reduction of plastid transcription, translation, and tetrapyrrole synthesis (Hanaoka *et al.* 2003, Woodson *et al.* 2013). However, the *sig2* mutation is not able to suppress the *fc2-1* phenotype as cell death was still observed in *fc2-1 sig2-2* mutants under diurnal cycling light conditions (Figs. 7 a, b, and c). As such, this result suggests that *ppr30* and *mterf9* mutations are blocking the accumulation of a specific component(s) of the mature chloroplast that must be both produced and in place for chloroplasts to respond to ^1^O_2_ accumulation. Such a switch may allow developing seedlings and leaves to establish mature photosynthesizing chloroplasts before they employ any chloroplast quality control pathways.

The role of such a pathway appears to be independent of the *fc2-1* mutation. When the *ppr30* and *mterf9* mutants were outcrossed to wt, they exhibited delayed photo-bleaching and cell death under a combination of moderate excess light and cold temperatures. Such stress has previously been shown to cause chloroplast ^1^O_2_ accumulation followed by cell death (Kim *et al.* 2012a, Meskauskiene *et al.* 2009, Triantaphylides *et al.* 2008). Therefore, the regulation of chloroplast gene expression may be one natural mechanism that allows photosynthetic eukaryotes to modulate growth and stress under dynamic and harsh environments. Support for this hypothesis has also been found in the green algae *Chlamydomonas reinhardtii.* The inhibition of plastid translation in this organism leads to a plastid unfolded protein response that induces nuclear genes involved in photo-oxidative stress (Perlaza *et al.* 2019). In plants, various *mterf* mutants have been shown to have altered stress responses (Leister *et al.* 2017). *mterf9* mutants have also been shown to have increased resistance to osmotic stress from salt and mannitol treatments (Robles *et al.* 2015). Loss of two other chloroplast-localized mTERF proteins, mTERF1 (Meskauskiene *et al.* 2009) and mTERF5 (Robles *et al.* 2012), lead to resistance to chloroplast ^1^O_2_, and osmotic stress, respectively. Finally, mutants lacking the mitochondrial localized mTERF4 (Quesada *et al.* 2011) and mTEF18 (Kim *et al.* 2012b) have increased and decreased tolerance to heat, respectively, suggesting that gene expression in the mitochondria can have similar effects on stress tolerance. Although the transcripts regulated by these mTERFs are unknown, together these reports suggest that organelle gene expression and stress are intimately linked in complex ways.

### Conclusions

The work presented here has demonstrated that full chloroplast gene expression, or the expression of a specific set of genes, is a prerequisite for initiating ^1^O_2_ signals from stressed chloroplasts. Future work will undoubtedly focus on identifying the specific targets of PPR30, mTERF9, and other proteins involved in post-transcriptional chloroplast gene expression. A deeper understanding of these proteins’ targets and their relation to stress should offer unique tools to engineer plants with specific responses to environmental stresses. Such knowledge of the control of photosynthesis and growth will aid our quest for abundant and inexpensive sources of food and fuel in the coming century.

## Experimental Procedures

### Biological material, growth conditions, and treatments

In this study, the wt line used in all experiments and to generate all transgenic constructs was *Arabidopsis thaliana* ecotype *Columbia* (Col-0). Table S2 lists the mutant lines used. Sequence data for the T-DNA lines came from the GABI (Kleinboelting *et al.* 2012), WiscDsLox (Woody *et al.* 2007), SAIL (Sessions *et al.* 2002), and SALK (The Salk Institute Genomic Analysis Laboratory) (Alonso *et al.* 2003) collections; *fc2-1* (GABI_766H08) (Woodson *et al.* 2011), *mterf9* (WiscDsLox474E07) (Robles *et al.* 2015), and *sig2-2* (Salk_045706) (Woodson *et al.* 2013) were described previously. Crossing was used to generate double mutant lines, which were confirmed by PCR-based markers listed in table S3.

Seeds were surface sterilized using 30% liquid bleach (v:v) with 0.04% Triton X-100 (v:v) for ten minutes. Seeds were pelleted at room temperature in a microfuge for 30 seconds at 2,000 × g. The seeds were then washed four times in sterile water and finally resuspended in 0.1% micropropagation type-1 agar powder (Caisson Laboratories). The seed suspension was then plated on Linsmaier and Skoog medium pH 5.7 (Caisson Laboratories North Logan, UT) with 0.6% micropropagation type-1 agar powder. A 3-4 day cold treatment of 4°C in the dark was used to stratify seeds before transferring them to constant light or diurnal light/dark cycling conditions of approximately ~80 μmol m^−2^ sec^−1^ white light at 22°C. For Pchlide measurement studies, germination was induced with only two hours of light exposure before the plates were wrapped in three layers of aluminum foil and incubated in the dark at 22°C. Seedlings were allowed to germinate and then harvested under dim green light for Pchlide extraction. For inducing excess light stress in adult plants, seedlings were grown on plates for 7 days in constant light conditions and then transferred to soil where they grew in 16h light/8h dark diurnal cycling light (~90-100 μmol m^−2^ sec^−1^ white light at 22°C) conditions. After 21 days in total, plants were moved under a LED panel (Hettich, Beverly, MA) and exposed to constant excess (650 μmol m^−2^ sec^−1^) or dim (6 μmol m^−2^ sec^−1)^ white light at 4°C. For transient expression experiments, *Nicotiana benthamiana* was grown under 16h light/8h dark diurnal cycling light conditions of ~100 μmol m^−2^ sec^−1^ white light at 22°C. Photosynthetically active radiation was measured using a LI-250A light meter with a LI-190R-BNC-2 Quantum Sensor (LiCOR). All bacteria (*E. coli* and *Agrobacterium tumefaciens* strains) were grown in liquid Miller nutrient broth or solid medium containing 1.5% agar (w/v). Cells were grown at 37°C (*E. coli*) or 28°C (*A. tumefaciens*) with the appropriate antibiotics and liquid medium was shaken at 225 rpm.

### Construction of complementation vectors

DNA fragments were amplified from wt *Arabidopsis* genomic DNA template using the Phusion enzyme (Finnzymes Espoo, Finland). These DNA fragments were gel-purified using the QIAquick Gel Extraction Kit (Qiagen) and cloned into the Gateway compatible vector pENTR-D/TOPO (Invitrogen) according to the manufacturer’s instructions. All primers and vectors are listed in tables S3 and S4, respectively.

DNA fragments were then transferred to either pEARLEYGATE101 (35S overexpression promoter plus C-terminal HA and YFP tags (Earley *et al.* 2006)) or pGBGWY (no promoter (Zhong *et al.* 2008)) to overexpress tagged YFP-HA proteins or to express native proteins with the native promoter, respectively. DNA fragments were then transferred to these vectors using LR clonase (Invitrogen) according to manufacturer’s instructions. The completed vectors were then transformed into the *A. tumefaciens* strain GV301, which was subsequently used to transform *Arabidopsis* via the floral dip method. T_1_ plants were selected for their ability to grow on Basta-soaked soil and propagated to the next generation. T_2_ lines were monitored for single insertions (segregating 3:1 for Basta resistance:sensitivity) and propagated to the next generation. Finally, homozygous lines were selected in the T_3_ generation based on 100% Basta-resistance

### RNA extraction and Real-time quantitative PCR

The RNeasy Plant Mini Kit (Qiagen) was used to extract total RNA from whole seedlings. Next, cDNA was synthesized using the Maxima first strand cDNA synthesis kit for RT-qPCR with DNase (Thermo Scientific) following the manufacturer’s instructions. Real-time PCR was performed using the SYBR Green Master Mix (BioRad) with the SYBR Green fluorophore and a CFX Connect Real Time PCR Detection System (Biorad). The following 2-step thermal profile was used in all RT-qPCR: 95 °C for 3 min, 40 cycles of 95 °C for 10s and 60 °C for 30s. *Actin2* expression was used as a standard to normalize all gene expression data. Table S3 lists the primers used.

### Chlorophyll measurements

Total chlorophyll was extracted from whole seedlings with 80% acetone and measured as previously described (Woodson *et al.* 2015). Levels were calculated spectrophotometrically according to (Hendry and Grime 1993).

### Protochlorophyllide measurements

Pchlide was extracted from whole seedlings with 80% acetone and measured by fluorescence using an adapted protocol (Shin *et al.* 2009) as previously described (Woodson *et al.* 2015).

### ALA measurements

ALA accumulation was measured as previously described (Woodson *et al.* 2015) using an adaptation of a previously described protocol (Czarnecki *et al.* 2011).

### Monitoring Singlet oxygen accumulation

To measure ^1^O_2_ accumulation in cotyledon mesophyll cells, we used a previously described method (Woodson *et al.* 2015) with minor alterations. Seedlings were grown on plates in 6h light/18h dark diurnal cycling light conditions. At the end of day 3, seedlings were transferred to 1.5 ml centrifuge tubes containing 250 μl of ½-strength Linsmaier liquid media and subsequently wrapped in foil and incubated in the dark at 22°C overnight. 60 min prior to dawn on the fourth day, 50 μM of 1.5 mM Singlet Oxygen Sensor Green (SOSG, Molecular Probes) and 0.1% Tween 20 (v/v) was added to the medium under dim light (final [250 μM]. Seedlings were vacuum infiltrated for 30 min and placed back in the dark. After 30 additional minutes, seedlings were exposed to light (dawn number four) for two hours. Seedlings were then washed twice with 1 ml of ½-strength Linsmaier and then imaged using a Zeiss Axiozoom 16 fluorescent stereo microscope equipped with a Hamamatsu Flash 4.0 camera and a GFP fluorescence filter. At least ten seedlings from each genotype were monitored and average fluorescence per mm^2^ was quantified using ImageJ.

### Trypan Blue staining

Trypan blue staining was used to assess cell death in leaves and cotyledons using a previously described method (Woodson *et al.* 2015). To quantify cell death, trypan blue intensity was measured with ImageJ using at least six seedlings from each genotype.

### In vivo protein localization

Expression constructs were transiently expressed in *N. benthamiana* leaves prior to imaging. To transform *N. benthamiana* leaves, dense 5 ml *A. tumefaciens* cultures harboring the appropriate vector were pelleted (2 minutes at 10,000 × g) and resuspended in 1 ml infiltration buffer (10 mM MES pH 5.6, 10 mM MgCl_2_). The optical density of the culture was adjusted to OD600 = 0.8, and 1.5 μl of Acetosyringone (100 mM stock in 100% DMSO) was added per 1 ml infiltration buffer. Cultures were incubated in the dark at room temperature for two hours before being infiltrated into leaves using a blunt end syringe. Tobacco plants were then grown normally and imaged three days later. Leaves were imaged using a Leica SP/2 inverted laser scanning confocal microscope. Image analysis was performed using the Leica SP/2 software package and the ImageJ (Schneider *et al.* 2012) bundle (Wright Cell Imaging facility).

### Electron Microscopy

Four day old seedlings were removed from LS plates and plunged into a solution of 2.5% glutaraldehyde and 2% formaldehyde (v:v) in 0.133 M Sorensen’s buffer (pH 7.4). Primary fixation was carried out under vacuum for approximately 18 hours. Seedlings were then washed three times with Sorenson’s buffer (10 minutes per wash) before being transferred to the secondary fixation solution of 1% osmium tetroxide and 1.5% potassium ferrocyanide in buffer for 90 minutes. Seedlings were then rinsed three times in ddH_2_0 (10 minutes per rinse) before *enbloc* staining in 2% uranyl acetate for 2 hours, followed again by three rinses in ddH_2_0. All fixations were carried out at room temperature. Seedlings were dehydrated in an ascending ethanol series (20%, 40%…) and infiltrated in 25% increments with Spurr’s resin (Spurr 1969). Infiltrated seedlings were placed in flat silicone molds with fresh Spurr’s resin before polymerization at 60°C for 24 hours. Selected seedlings were sectioned on a Reichert UCT ultramicrotome (Leica Microsystems Inc., Bannockburn, IL), collected on copper grids, and imaged with a FEI Tecnai Spirit Transmission Electron Microscope at 100kV (ThermoFisher, Waltham, MA).

Chloroplast area and chloroplast compartment size (% chloroplast area/total cell area) were analyzed using ImageJ. Average chloroplast size was calculated on a per cell basis. For chloroplast compartment ratios, the sum of all chloroplast areas was divided by the total cell area. Only intact and fully imaged cells were used for these calculations. Measurements were collected from at least three cells in three different seedlings in each genotype and growth condition.

### Statistical analyses

All reported statistical analyses (one-way ANOVA and post-tests (Tukey-Kramer and Dunnett’s multiple comparisons)) were performed using GraphPad Prism software.

## Acknowledgements

We thank Dr. Alice Barkan (U. Oregon) for useful discussions about PPR30. The authors acknowledge the Division of Chemical Sciences, Geosciences, and Biosciences, Office of Basic Energy Sciences of the U.S. Department of Energy grants DE-SC0019573 and DE-FG02-04ER15540 awarded to J.D.W. and J.C., respectively. The authors have no conflict of interest to declare.

## Short legends for supporting information

### Supplemental Tables

Table S1. Location of confirmed *ferrochelatase two suppressor* (*fts*) mutations

Table S2. Mutant plant lines used in study.

Table S3. Primers used in study

Table S4. Vectors used in study

### Supplemental Figures

Figure S1. Mapping of *fts3*, *fts32*, and *fts38*.

Figure S2. PPR30 and mTERF9 are localized to the chloroplasts.

Figure S3. *fts32* and *fts38* have delayed chloroplast development.

Figure S4. *ppr30* and *mTERF9* mutants are resistant to excess light stress.

## Data Statement

All data used in the manuscript are included within. All biological materials generated for this work will be available upon request.

## Notes

#### Summary of Updates

The introduction, results, and discussion sections have been extended for clarification and to include additional background information. The data, figures, findings, and conclusions remain the same.

## References

Alonso, J.M., Stepanova, A.N., Leisse, T.J., Kim, C.J., Chen, H., Shinn, P., Stevenson, D.K., Zimmerman, J., Barajas, P., Cheuk, R., Gadrinab, C., Heller, C., Jeske, A., Koesema, E., Meyers, C.C., Parker, H., Prednis, L., Ansari, Y., Choy, N., Deen, H., Geralt, M., Hazari, N., Hom, E., Karnes, M., Mulholland, C., Ndubaku, R., Schmidt, I., Guzman, P., Aguilar-Henonin, L., Schmid, M., Weigel, D., Carter, D.E., Marchand, T., Risseeuw, E., Brogden, D., Zeko, A., Crosby, W.L., Berry, C.C. and Ecker, J.R. (2003) Genome-wide insertional mutagenesis of Arabidopsis thaliana. Science, 301, 653–657.

Asada, K. (2006) Production and scavenging of reactive oxygen species in chloroplasts and their functions. Plant Physiol, 141, 391–396.

Babiychuk, E., Vandepoele, K., Wissing, J., Garcia-Diaz, M., De Rycke, R., Akbari, H., Joubes, J., Beeckman, T., Jansch, L., Frentzen, M., Van Montagu, M.C. and Kushnir, S. (2011) Plastid gene expression and plant development require a plastidic protein of the mitochondrial transcription termination factor family. Proc Natl Acad Sci U S A, 108, 6674–6679.

Barkan, A. and Goldschmidt-Clermont, M. (2000) Participation of nuclear genes in chloroplast gene expression. Biochimie, 82, 559–572.

Barkan, A. and Small, I. (2014) Pentatricopeptide repeat proteins in plants. Annu Rev Plant Biol,65, 415–442.

Baruah, A., Simkova, K., Apel, K. and Laloi, C. (2009) Arabidopsis mutants reveal multiple singlet oxygen signaling pathways involved in stress response and development. Plant Mol Biol, 70, 547–563.

Chan, K.X., Mabbitt, P.D., Phua, S.Y., Mueller, J.W., Nisar, N., Gigolashvili, T., Stroeher, E., Grassl, J., Arlt, W., Estavillo, G.M., Jackson, C.J. and Pogson, B.J. (2016) Sensing and signaling of oxidative stress in chloroplasts by inactivation of the SAL1 phosphoadenosine phosphatase. Proc Natl Acad Sci U S A, 113, E4567–4576.

Chan, K.X., Phua, S.Y., Crisp, P., McQuinn, R. and Pogson, B.J. (2015) Learning the Languages of the Chloroplast: Retrograde Signaling and Beyond. Annu Rev Plant Biol, 67, 25–53.

Coll, N.S., Danon, A., Meurer, J., Cho, W.K. and Apel, K. (2009) Characterization of soldat8, a suppressor of singlet oxygen-induced cell death in Arabidopsis seedlings. Plant Cell Physiol, 50, 707–718.

Czarnecki, O., Peter, E. and Grimm, B. (2011) Methods for analysis of photosynthetic pigments and steady-state levels of intermediates of tetrapyrrole biosynthesis. Methods Mol Biol, 775, 357–385.

Dogra, V., Duan, J., Lee, K.P., Lv, S., Liu, R. and Kim, C. (2017) FtsH2-Dependent Proteolysis of EXECUTER1 Is Essential in Mediating Singlet Oxygen-Triggered Retrograde Signaling in Arabidopsis thaliana. Front Plant Sci, 8, 1145.

Dogra, V., Li, M., Singh, S., Li, M. and Kim, C. (2019) Oxidative post-translational modification of EXECUTER1 is required for singlet oxygen sensing in plastids. Nat Commun, 10, 2834.

Doniwa, Y., Ueda, M., Ueta, M., Wada, A., Kadowaki, K. and Tsutsumi, N. (2010) The involvement of a PPR protein of the P subfamily in partial RNA editing of an Arabidopsis mitochondrial transcript. Gene, 454, 39–46.

Earley, K.W., Haag, J.R., Pontes, O., Opper, K., Juehne, T., Song, K. and Pikaard, C.S. (2006) Gateway-compatible vectors for plant functional genomics and proteomics. Plant J, 45, 616–629.

Foyer, C.H.. (2018) Reactive oxygen species, oxidative signaling and the regulation of photosynthesis. Environ Exp Bot, 154, 134–142.

Gutierrez, J., Gonzalez-Perez, S., Garcia-Garcia, F., Daly, C.T., Lorenzo, O., Revuelta, J.L., McCabe, P.F. and Arellano, J.B. (2014) Programmed cell death activated by Rose Bengal in Arabidopsis thaliana cell suspension cultures requires functional chloroplasts. J Exp Bot, 65, 3081–3095.

Hanaoka, M., Kanamaru, K., Takahashi, H. and Tanaka, K. (2003) Molecular genetic analysis of chloroplast gene promoters dependent on SIG2, a nucleus-encoded sigma factor for the plastid-encoded RNA polymerase, in Arabidopsis thaliana. Nucleic Acids Res, 31, 7090–7098.

Hendry, G. and Grime, J.P. (1993) Methods in Comparative Plant Ecology - A Laboratory Manual. 148–152.

Jarvis, P. and Lopez-Juez, E. (2013) Biogenesis and homeostasis of chloroplasts and other plastids. Nat Rev Mol Cell Biol, 14, 787–802.

Kim, C., Meskauskiene, R., Zhang, S., Lee, K.P., Lakshmanan Ashok, M., Blajecka, K., Herrfurth, C., Feussner, I. and Apel, K. (2012a) Chloroplasts of Arabidopsis are the source and a primary target of a plant-specific programmed cell death signaling pathway. Plant Cell, 24, 3026–3039.

Kim, M., Lee, U., Small, I., des Francs-Small, C.C. and Vierling, E. (2012b) Mutations in an Arabidopsis mitochondrial transcription termination factor-related protein enhance thermotolerance in the absence of the major molecular chaperone HSP101. Plant Cell, 24, 3349–3365.

Kleinboelting, N., Huep, G., Kloetgen, A., Viehoever, P. and Weisshaar, B. (2012) GABI-Kat SimpleSearch: new features of the Arabidopsis thaliana T-DNA mutant database. Nucleic Acids Res, 40, D1211–1215.

Kleine, T. (2012) Arabidopsis thaliana mTERF proteins: evolution and functional classification. Front Plant Sci, 3, 233.

Koussevitzky, S., Nott, A., Mockler, T.C., Hong, F., Sachetto-Martins, G., Surpin, M., Lim, J., Mittler, R. and Chory, J. (2007) Signals from chloroplasts converge to regulate nuclear gene expression. Science, 316, 715–719.

Leister, D., Wang, L. and Kleine, T. (2017) Organellar Gene Expression and Acclimation of Plants to Environmental Stress. Front Plant Sci, 8, 387.

Li, P., Ponnala, L., Gandotra, N., Wang, L., Si, Y., Tausta, S.L., Kebrom, T.H., Provart, N., Patel, R., Myers, C.R., Reidel, E.J., Turgeon, R., Liu, P., Sun, Q., Nelson, T. and Brutnell, T.P. (2010) The developmental dynamics of the maize leaf transcriptome. Nat Genet, 42, 1060–1067.

Lu, Y. and Yao, J. (2018) Chloroplasts at the Crossroad of Photosynthesis, Pathogen Infection and Plant Defense. Int J Mol Sci, 19.

Majeran, W., Friso, G., Asakura, Y., Qu, X., Huang, M., Ponnala, L., Watkins, K.P., Barkan, A. and van Wijk, K.J. (2012) Nucleoid-enriched proteomes in developing plastids and chloroplasts from maize leaves: a new conceptual framework for nucleoid functions. Plant Physiol, 158, 156–189.

Meskauskiene, R., Wursch, M., Laloi, C., Vidi, P.A., Coll, N.S., Kessler, F., Baruah, A., Kim, C. and Apel, K. (2009) A mutation in the Arabidopsis mTERF-related plastid protein SOLDAT10 activates retrograde signaling and suppresses (1)O(2)-induced cell death. Plant J, 60, 399–410.

Ogilby, P.R. (2010) Singlet oxygen: there is indeed something new under the sun. Chemical Society reviews, 39, 3181–3209.

op den Camp, R.G., Przybyla, D., Ochsenbein, C., Laloi, C., Kim, C., Danon, A., Wagner, D., Hideg, E., Gobel, C., Feussner, I., Nater, M. and Apel, K. (2003) Rapid induction of distinct stress responses after the release of singlet oxygen in Arabidopsis. Plant Cell, 15, 2320–2332.

Perlaza, K., Toutkoushian, H., Boone, M., Lam, M., Iwai, M., Jonikas, M.C., Walter, P. and Ramundo, S. (2019) The Mars1 kinase confers photoprotection through signaling in the chloroplast unfolded protein response. Elife, 8.

Pesaresi, P. and Kim, C. (2019) Current understanding of GUN1: a key mediator involved in biogenic retrograde signaling. Plant Cell Rep, 38, 819–823.

Quesada, V., Sarmiento-Manus, R., Gonzalez-Bayon, R., Hricova, A., Perez-Marcos, R., Gracia-Martinez, E., Medina-Ruiz, L., Leyva-Diaz, E., Ponce, M.R. and Micol, J.L. (2011) Arabidopsis RUGOSA2 encodes an mTERF family member required for mitochondrion, chloroplast and leaf development. Plant J, 68, 738–753.

Ramel, F., Ksas, B., Akkari, E., Mialoundama, A.S., Monnet, F., Krieger-Liszkay, A., Ravanat, J.L., Mueller, M.J., Bouvier, F. and Havaux, M. (2013) Light-induced acclimation of the Arabidopsis chlorina1 mutant to singlet oxygen. Plant Cell, 25, 1445–1462.

Robles, P., Micol, J.L. and Quesada, V. (2012) Arabidopsis MDA1, a nuclear-encoded protein, functions in chloroplast development and abiotic stress responses. PLoS One, 7, e42924.

Robles, P., Micol, J.L. and Quesada, V. (2015) Mutations in the plant-conserved MTERF9 alter chloroplast gene expression, development and tolerance to abiotic stress in Arabidopsis thaliana. Physiol Plant, 154, 297–313.

Savage, L.J., Imre, K.M., Hall, D.A. and Last, R.L. (2013) Analysis of essential Arabidopsis nuclear genes encoding plastid-targeted proteins. PLoS One, 8, e73291.

Scharfenberg, M., Mittermayr, L., E, V.O.N.R.-L., Schlicke, H., Grimm, B., Leister, D. and Kleine, T. (2015) Functional characterization of the two ferrochelatases in Arabidopsis thaliana. Plant Cell Environ, 38, 280–298.

Schneeberger, K., Ossowski, S., Lanz, C., Juul, T., Petersen, A.H., Nielsen, K.L., Jorgensen, J.E., Weigel, D. and Andersen, S.U. (2009) SHOREmap: simultaneous mapping and mutation identification by deep sequencing. Nat Methods, 6, 550–551.

Schneider, C.A., Rasband, W.S. and Eliceiri, K.W. (2012) NIH Image to ImageJ: 25 years of image analysis. Nat Methods, 9, 671–675.

Sessions, A., Burke, E., Presting, G., Aux, G., McElver, J., Patton, D., Dietrich, B., Ho, P., Bacwaden, J., Ko, C., Clarke, J.D., Cotton, D., Bullis, D., Snell, J., Miguel, T., Hutchison, D., Kimmerly, B., Mitzel, T., Katagiri, F., Glazebrook, J., Law, M. and Goff, S.A. (2002) A high-throughput Arabidopsis reverse genetics system. Plant Cell, 14, 2985–2994.

Shimizu, T., Kacprzak, S.M., Mochizuki, N., Nagatani, A., Watanabe, S., Shimada, T., Tanaka, K., Hayashi, Y., Arai, M., Leister, D., Okamoto, H., Terry, M.J. and Masuda, T. (2019) The retrograde signaling protein GUN1 regulates tetrapyrrole biosynthesis. Proc Natl Acad Sci U S A.

Shin, J., Kim, K., Kang, H., Zulfugarov, I.S., Bae, G., Lee, C.H., Lee, D. and Choi, G. (2009) Phytochromes promote seedling light responses by inhibiting four negatively-acting phytochrome-interacting factors. Proc Natl Acad Sci U S A, 106, 7660–7665.

Small, I.D. and Peeters, N. (2000) The PPR motif - a TPR-related motif prevalent in plant organellar proteins. Trends Biochem Sci, 25, 46–47.

Spurr, A.R. (1969) A low-viscosity epoxy resin embedding medium for electron microscopy. J Ultrastruct Res, 26, 31–43.

Suo, J., Zhao, Q., David, L., Chen, S. and Dai, S. (2017) Salinity Response in Chloroplasts: Insights from Gene Characterization. Int J Mol Sci, 18.

Tanaka, R., Kobayashi, K. and Masuda, T. (2011) Tetrapyrrole Metabolism in Arabidopsis thaliana. Arabidopsis Book, 9, e0145.

Triantaphylides, C., Krischke, M., Hoeberichts, F.A., Ksas, B., Gresser, G., Havaux, M., Van Breusegem, F. and Mueller, M.J. (2008) Singlet oxygen is the major reactive oxygen species involved in photooxidative damage to plants. Plant Physiol, 148, 960–968.

van Wijk, K.J. and Baginsky, S. (2011) Plastid proteomics in higher plants: current state and future goals. Plant Physiol, 155, 1578–1588.

Wagner, D., Przybyla, D., Op den Camp, R., Kim, C., Landgraf, F., Lee, K.P., Wursch, M., Laloi, C., Nater, M., Hideg, E. and Apel, K. (2004) The genetic basis of singlet oxygen-induced stress responses of Arabidopsis thaliana. Science, 306, 1183–1185.

Woodson, J.D. (2016) Chloroplast quality control - balancing energy production and stress. New Phytol, 212, 36–41.

Woodson, J.D. (2019) Chloroplast stress signals: regulation of cellular degradation and chloroplast turnover. Curr Opin Plant Biol, 52, 30–37.

Woodson, J.D., Joens, M.S., Sinson, A.B., Gilkerson, J., Salome, P.A., Weigel, D., Fitzpatrick, J.A. and Chory, J. (2015) Ubiquitin facilitates a quality-control pathway that removes damaged chloroplasts. Science, 350, 450–454.

Woodson, J.D., Perez-Ruiz, J.M. and Chory, J. (2011) Heme synthesis by plastid ferrochelatase I regulates nuclear gene expression in plants. Curr Biol, 21, 897–903.

Woodson, J.D., Perez-Ruiz, J.M., Schmitz, R.J., Ecker, J.R. and Chory, J. (2013) Sigma factor-mediated plastid retrograde signals control nuclear gene expression. Plant J, 73, 1–13.

Woody, S.T., Austin-Phillips, S., Amasino, R.M. and Krysan, P.J. (2007) The WiscDsLox T-DNA collection: an arabidopsis community resource generated by using an improved high-throughput T-DNA sequencing pipeline. J Plant Res, 120, 157–165.

Wu, G.Z., Meyer, E.H., Richter, A.S., Schuster, M., Ling, Q., Schottler, M.A., Walther, D., Zoschke, R., Grimm, B., Jarvis, R.P. and Bock, R. (2019) Control of retrograde signalling by protein import and cytosolic folding stress. Nat Plants, 5, 525–538.

Yagi, Y. and Shiina, T. (2014) Recent advances in the study of chloroplast gene expression and its evolution. Front Plant Sci, 5, 61.

Yu, Q.B., Huang, C. and Yang, Z.N. (2014) Nuclear-encoded factors associated with the chloroplast transcription machinery of higher plants. Front Plant Sci, 5, 316.

Zhao, X., Huang, J. and Chory, J. (2019) GUN1 interacts with MORF2 to regulate plastid RNA editing during retrograde signaling. Proc Natl Acad Sci U S A, 116, 10162–10167.

Zhong, S., Lin, Z., Fray, R.G. and Grierson, D. (2008) Improved plant transformation vectors for fluorescent protein tagging. Transgenic Res, 17, 985–989.

