## Supporting information for "Plastid gene expression is required for singlet oxygen-induced chloroplast degradation and cell death"

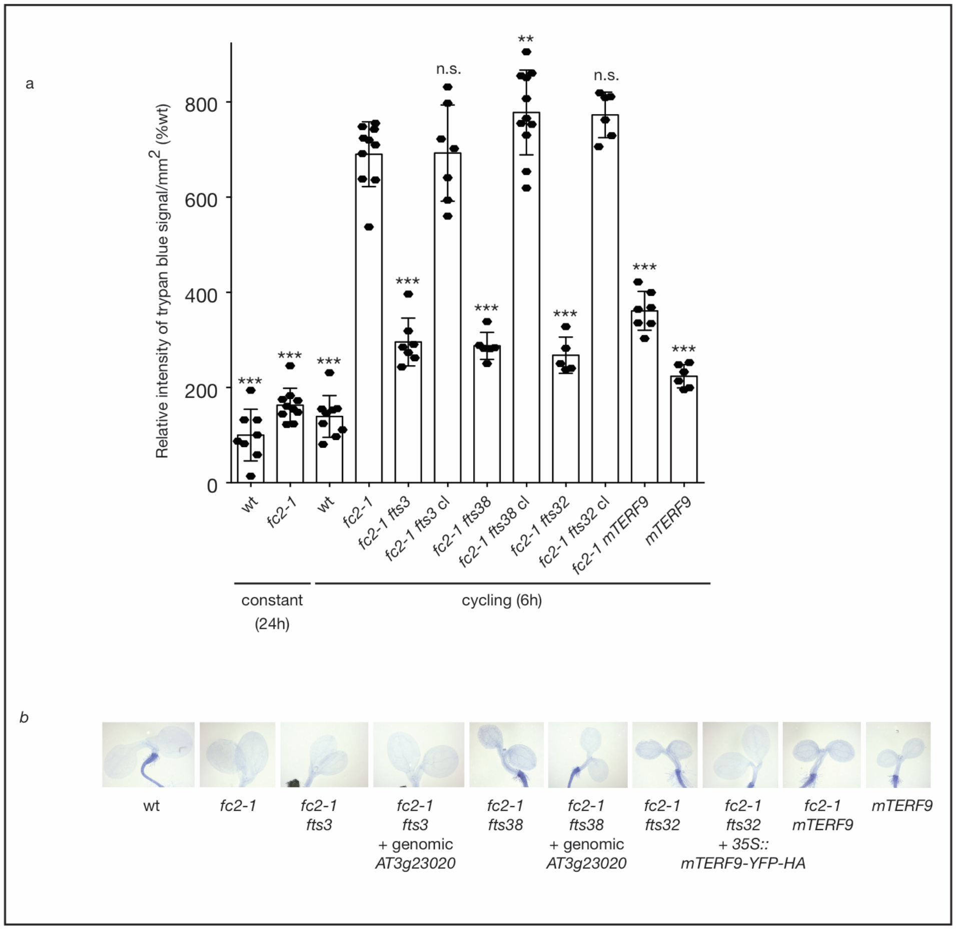


Figure S1

Figure S1. Mapping of *fts3*, *fts32*, and *fts38*. **A)** Quantification of trypan blue staining (relative intensity/mm$^2$) of six-day-old seedlings grown in constant (24h) light or 6h light / 18h dark diurnal cycling conditions. Shown are means of biological replicates indicted by closed circles ($n \geq 5$). Statistical analyses were performed by a one-way ANOVA followed by Dunnett's multiple comparisons test with the *fc2-1* in cycling light sample. **, ***, and n.s. indicates an adjusted p value of $\leq 0.01$, $\leq 0.001$, and $\geq 0.05$, respectively. **B)** Shown are representative six-day-old seedlings grown in constant light and stained with trypan blue. The absence of any deep dark color is indicative of no cell death.

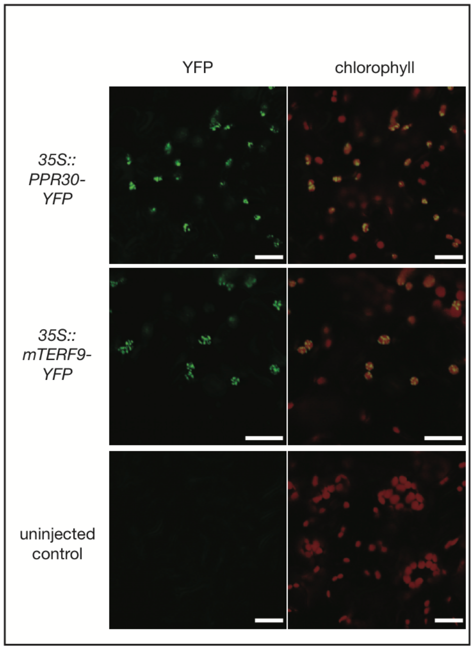


Figure S2

Figure S2. PPR30 and mTERF9 are localized to the chloroplasts.

Shown are representative confocal microscopy images of transiently expressed PPR30- and mTERF9-YFP protein co-localizing with chlorophyll auto-fluorescence in *Nicotiana benthamiana* leaf mesophyll cells. Scale bars = 20 µm.

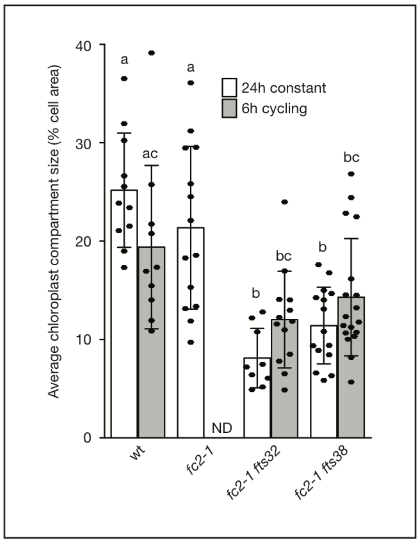


Figure S3

Figure S3. *fts32* and *fts38* have delayed chloroplast development. Shown are means (+/- SD) of the average % chloroplast compartment size (total chloroplast area/cell area) of four day old seedlings grown in constant (24h) light or 6h light / 18h dark diurnal cycling conditions as assessed by TEM (n ≥ 9 cells). Individual measurements are indicated by closed circles. Chloroplast volume in *fc2-1* seedlings grown in cycling light conditions was not determined due to extensive cellular degradation. The chlorophyll statistical analysis was performed by a one-way ANOVA and different letters above bars indicate significant differences determined by a Tukey's test (p value ≤ 0.05).

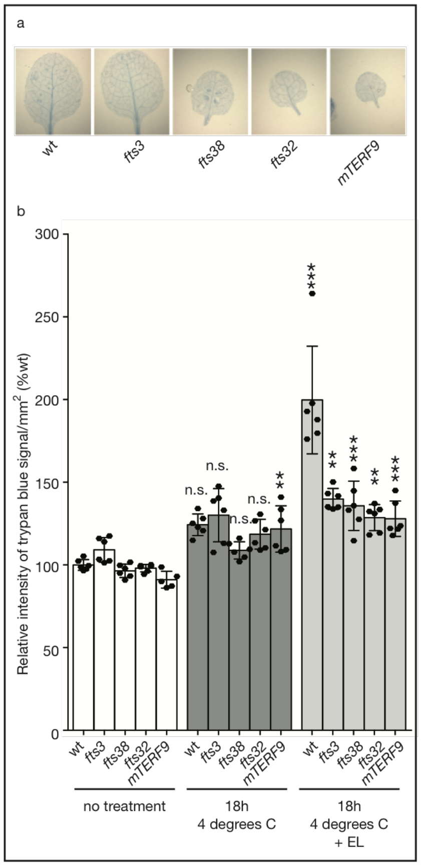


Figure S4

Figure S4. *ppr30* and *mTERF9* mutants are resistant to excess light stress.

**A)** Shown are representative images of leaves exposed to 18 hours 4°C in dim light (~ 6 μmol $m^{-2}$ $sec^{-1}$) and then stained by trypan blue. The lack of any deep blue color is indicative of healthy tissue with no cell death. **B)** Shown are the means of trypan blue staining of single leaves exposed to 18 hours 4°C plus dim light or 4°C plus excess light (n = 6 leaves per genotype per treatment). Individual measurements are indicated by closed circles. Statistical analysis was performed by a one-way ANOVA followed by Dunnett's multiple comparisons test with the control sample (no 4°C or excess light treatment) of each genotype. *, ***, and n.s. indicates an adjusted p value of ≤ 0.05, ≤ 0.001, and ≥ 0.05, respectively.

**Table S1.**

Location of confirmed *ferrochelatase two suppressor* (*fts*) mutations

| Mutant | Gene name | Gene # | mutation | Change in protein sequence |
|---|---|---|---|---|
| *fts3/ppr30-1* | *PPR30* | At3g23020 | c8177451t | D765N |
| *fts32/mterf9-3* | *mTERF9/mTERF31* | At5g55580 | c22516932t | R368stop |
| *fts38/ppr30-2* | *PPR30* | At3g23020 | c8177283t | G821R |

**Table S2.**

Mutant plant lines used in study

| Mutant | Gene | mutation | notes | ref |
|---|---|---|---|---|
| *fc2-1* | *Plastid Ferrochelatase 2, FC2, At2g30390* | GABI_766H08 T-DNA in 5'UTR | Sulfadiazin$^r$ | (Woodson *et al.* 2011) |
| *mterf9* | *Mitochondrial Transcription Termination Factor 9 (mTERF9), At5g55580* | WiscDsLox474E07, T-DNA in exon 4/4 | | (Robles *et al.* 2015) |
| *sig2-2* | *SIG2 / At1g08540* | Salk_045706, T-DNA in Intron 1/6 | | (Woodson *et al.* 2013) |
| *gun1-9* | *AT2G31400 / Genomes Uncoupled 1* | Single point mutation leading to premature stop codon (Q654Stop) | | (Koussevitzky *et al.* 2007) |

**Table S3.**

Primers used in study

| **Gene** | **Primer orientation / name** | **Sequence** |
|---|---|---|
| **qPCR primer pairs** | | |
| *ACTIN2 At3g18780* | For. / JP199 | GCACTTGCACCAAGCAGCAT |
| | Rev. / JP200 | CCTTTCAGGTGGTGCAACGAC |
| *LHCB 1.2, CAB3 At1g29910* | For. / JP197 | GGACTTGCTTTACCCCGGTG |
| | Rev. / JP198 | TCGGTAGCAAGACCCAATGG |
| *LHCB 2.2, At2g05070* | Rev. / WLO1529 | GCTTTGTAAACTCGTGATTGTG |
| | For. / WLO1530 | TGCCAAATTCACATCAAACG |
| *RBCS 2B, At5G38420* | Rev. / WLO1448 | GCTTCACCGAAGCTTAATCC |
| | For. / WLO1449 | CCACATAGAAATGGGTTCCAG |
| *CA1 AT3g01500* | For. / JP209 | TGTGTCCATCACACGTTCTGG |
| | Rev. / JP210 | GGACCACGAAGGCATCTCCT |
| *psaJ AtCG00630* | Rev. / WLO1539 | GTACTCTATGGTTCGGTTCGT |
| | For. / WLO1540 | AGGGAAATGTTAATGCATCTGG |
| *psbA AtCG00020* | For. / WLO1535 | CGTCTTTACATTGGATGGTTTGG |
| | Rev. / WLO1536 | CAGAAGTTGCGGTCAATAAGG |
| *psbB AtCG00680* | For. / WLO1537 | TCGTGCGACTTTGAAATCTGA |
| | Rev. / WLO1538 | CAACCTCTTGGGCTGCTACG |
| *rbcL AtCG00490* | For. / WLO1541 | TTACAAAGGACGATGCTACCACAT |
| | Rev. / WLO1542 | TGAGTTTCTTCTCCTGGAACGG |
| *clpP AtCG00670* | For. / WLO1545 | TGGGTTGACATATACAACCGACTTT |
| | Rev. / WLO1546 | GCCTAAAAAAAATAATCTTTCTCGATAAA |
| *rpoB AtCG00190* | For. / WLO1547 | ATACGAGATATCCATCCTAGTCAC |
| | Rev. / WLO1548 | GTCCAACATTGATTCCTTCAGAC |
| *trnEYD ATCG00250-230* | For. / WLO1559 | TCTAGTGGTTCAGGACATCTC |
| | Rev. / WLO1560 | TTGCCAACGAATTTACAGTC |
| *SIB1 At3g56710* | For. / JP589 | CAACCGGAGCCCATCTATT |
| | Rev. / JP590 | GGAGAAAGGTTGTGGTCGTC |
| *HSP26.5 At1g52560* | For. / JP585 | CGAGCTTATCGTTGCCTGAT |
| | Rev. / JP586 | CTCCGCCTTAATGTCCTCAA |
| *BAP1* At3g61190 | For. / JP338 | ATTGATGGATACGGTGGCCG |
| | Rev. / JP339 | CAGACCCCAAACCGGAACTC |
| *Atpase* At3g28580 | For. / JP336 | GAAGATCGGAAAAGCGTGGAA |
| | Rev. / JP337 | CCGGGTGGTCCAAACAAAAG |
| *ZAT12* At5g59820 | For. / JP344 | GCGTTGGTTACACGCGCTT |
| | Rev. / JP345 | CTTCAACGTAGTCACCGTGGG |
| *CYC8* At4g37370 | For. / JP1130 | AATGGGCATTGTCGAACGTG |
| | Rev. / JP1131 | TCGCCTTGTTCAATACATCCG |
| **Genotyping primers** | | |
| SALK T-DNA left border | LB1.3 | ATTTTGCCGATTTCGGAAC |

| | | |
|---|---|---|
| WiscDsLox T-DNA left border | p745 WISC T-DNA LB | AACGTCCGCAATGTGTTATTAAGTTGTC |
| SAIL T-DNA left border | LB1 | GCCTTTTCAGAAATGGATAAATAGCCTTG CTTCC |
| GABI-KAT Right border | Gabi-KAT 03144 | GTGGATTGATGTGATATCTCC |
| GABI-KAT Left border | Gabi-KAT 08409 | ATATTGACCATCATACTCATTGC |
| *fc2-1* (GabiKat_766H08) | LP / JP283 | GAGCAACGCCAAACATAGAAG |
| | RP / JP284 | TCAAAGGCAATGAATGTTTCC |
| *mterf9* (WiscDsLox474E07) | LP / 816 | CAAAACCTGGAAAAGATTGAGG |
| | RP / 817 | GGTCTTGGCATTCCTAATTCC |
| *sig2-2* (Salk_045706) | LP / JP39 | CTTCTTCGTCTTCATCATCCG |
| | RP / JP40 | CATTGAAGATTTGGGAACCAC |
| *fts3/ppr30-1* | LP / JP654 | TCGACATCCCAAGTTTCATCAG |
| | RP / JP655 | CGTTCTCGGCTTATTCGCTCTA (*Xba*I restriction enzyme cuts wt sequence) |
| *fts32/ mterf9-3* | LP / JP668 | CGACATAACATGGAACACTC |
| | RP / JP669 | TGCTCCTAAGGAAATTGACTC (*Xho*I restriction enzyme cuts wt sequence) |
| *fts38/ppr30-2* | LP / JP652 | GTTGAGATCCATAGCTCTAAGC |
| | RP / JP653 | TCAACCTGATGACTCAACGTTC (*HinD*III restriction enzyme cuts mutant sequence) |
| **Cloning primers** | | |
| At3g23020/PPR30 coding region for Gateway-no stop codon (2,548 bp fragment) | For / JP1061 | CACCGCTCGTTGCAGTTGAAAAATGT |
| | Rev / JP1062 | AAGCTCGTCGACACAATCTCC |
| At3g23020/PPR30 promoter and coding region for Gateway (5,415 bp fragment including 1.9 kb 5’ / 1 kb 3’ of coding region) | Rev / AS49 | GCGAACTGCGTCTTCTGAAC |
| | For / AS50 | CACCACCCGGAAGAACAGAGAAGA |
| At5g55580/mTERF9 coding region for Gateway-no stop codon (1,809 bp fragment) | For / JP1037 | CACCATGGCGGGTTTCTCACTGTAC |
| | Rev / JP1038 | TCCTCTCTTGTCATACTTGTTTGC |

**Table S4.**
Vectors used in study

| Plasmid | Insert | notes | ref |
|---|---|---|---|
| pENTR-D/TOPO | | Gateway cloning vector | Invitrogen |
| PEARLEYGATE-101 | | 35S promoter, C-term YFP-HA fusion | (Earley *et al.* 2006) |
| pGBGWY | | No promoter, C-term YFP | (Zhong *et al.* 2008) |
| pJDW199 | *FTS30 – stop codon* | pENTR-D/TOPO | This study |
| pJDW188 | *MTERF9 – stop codon* | pENTR-D/TOPO | This study |
| pABS11 | *Genomic fragment of FTS30* | pENTR-D/TOPO | This study |
| pJDW200 | *35S::FTS30-YFP-HA* | In PEARLEYGATE-101 | This study |
| pJDW206 | *35S::mTERF9-YFP-HA* | In PEARLEYGATE-101 | This study |
| pABS39 | *Genomic fragment of FTS30* | pGBGWY | This study |


### References

**Earley, K.W., Haag, J.R., Pontes, O., Opper, K., Juehne, T., Song, K. and Pikaard, C.S.** (2006) Gateway-compatible vectors for plant functional genomics and proteomics. *Plant J*, **45**, 616-629.

**Koussevitzky, S., Nott, A., Mockler, T.C., Hong, F., Sachetto-Martins, G., Surpin, M., Lim, J., Mittler, R. and Chory, J.** (2007) Signals from chloroplasts converge to regulate nuclear gene expression. *Science*, **316**, 715-719.

**Robles, P., Micol, J.L. and Quesada, V.** (2015) Mutations in the plant-conserved MTERF9 alter chloroplast gene expression, development and tolerance to abiotic stress in Arabidopsis thaliana. *Physiol Plant*, **154**, 297-313.

**Woodson, J.D., Perez-Ruiz, J.M. and Chory, J.** (2011) Heme synthesis by plastid ferrochelatase I regulates nuclear gene expression in plants. *Curr Biol*, **21**, 897-903.

**Woodson, J.D., Perez-Ruiz, J.M., Schmitz, R.J., Ecker, J.R. and Chory, J.** (2013) Sigma factor-mediated plastid retrograde signals control nuclear gene expression. *Plant J*, **73**, 1-13.

**Zhong, S., Lin, Z., Fray, R.G. and Grierson, D.** (2008) Improved plant transformation vectors for fluorescent protein tagging. *Transgenic Res*, **17**, 985-989.
